# Characterization of BoHV-4 ORF45

**DOI:** 10.1101/2022.12.07.519449

**Authors:** Luca Russo, Emanuele Capra, Valentina Franceschi, Davide Cavazzini, Roberto Sala, Barbara Lazzari, Sandro Cavirani, Gaetano Donofrio

**Affiliations:** Dipartimento di Scienze Medico Veterinarie, Università di Parma, Parma, Italy; Istituto di Biologia e Biotecnologia Agraria, Consiglio Nazionale delle Ricerche IBBA CNR, Lodi, Italy; Dipartimento di Scienze Chimiche, della Vita e della Sostenibilità Ambientale, Università di Parma, Parma, Italy; Dipartimento di Medicina e Chirurgia, Università di Parma, Parma, Italy

## Abstract

Bovine herpesvirus 4 (BoHV-4) is a *Gammaherpesvirus* of the genus *Rhadinovirus*, his natural host is the bovine whereas the African Buffalo the natural reservoir. Anyhow, BoHV-4 infection is not associated to a specific disease. Genome structure and genes are well conserved in *Gammaherpesvirus*, and *orf*45 gene and its product, ORF45, is one of those. BoHV-4 ORF45 has been suggested to be a tegument protein, however, BoHV-4 ORF45 structure and function have not yet been experimentally characterized. In the present study, it is shown that BoHV-4 ORF45, despite its poor homology with other characterized *Rhadinovirus* ORF45s, is structurally related to Kaposi’s sarcoma-associated herpesvirus (KSHV), is a phosphoprotein and localizes in the host cell nuclei. Through the generation of an ORF45-null mutant BoHV-4 and its pararevertant, it was possible to demonstrate that ORF45 is essential for BoHV-4 lytic replication and is associated to the viral particles, as for the other characterized *Rhadinovirus* ORF45s. Finally, the impact of BoHV-4 ORF45 on cellular transcriptome was investigated, an aspect poorly explored or not at all for other *Gammaherpesvirus*. Many cellular transcriptional pathways were found to be alterate, mainly those involving p90 ribosomal S6 kinase (RSK) and signal-regulated kinase (ERK) complex (RSK/ERK), thus highlighting the authentic character of BoHV-4 ORF45 and paving the way to further investigations.

## Introduction

Bovine herpesvirus 4 (BoHV-4) is a herpesvirus belonging to the *Gammaherpesvirinae* subfamily and *Rhadinovirus* genus. In some circumstances BoHV-4 was isolated from cattle affected by different pathological entities, such as abortion, metritis, pneumonia, diarrhea, respiratory infection, and mammary pustular dermatitis. However, the pathogenic role of BoHV-4 remains unclear and only a few investigators have successfully produced an experimental disease. Although BoHV-4 natural host is the bovine, in which was found to be widespread in the population and persists in many individuals as an asymptomatic infection, BoHV-4 was isolated from other ruminant species too(Todd and Storz, 1983; Wellemans et al., 1986; Moreno-Lopez et al., 1989) and occasionally from lions, cats, and owl monkeys (Barahona et al., 1973; Bublot et al., 1991). Worthy of note, BoHV-4 was found to be highly prevalent (higher than 68%) in African buffalo (*Syncerus Caffer*), the buffalo strain diverged from the taurine one 730,000 years ago (Dewals et al., 2006) and thus African Buffalo considered to be the natural reservoir of the virus (Dewals et al., 2005). Experimentally, BoHV-4 was also shown to infect goats (Moreno-Lopez et al., 1989), guinea pigs, and rabbits (Egyed et al., 1997). *In vitro*, BoHV-4 can replicate in primary cell cultures or cell lines from several host species, such as sheep, goats, swine, cats, dogs, rabbits, mink, horses, turkeys, ferrets, chickens, hamsters, rats, mice, monkeys, and human (Donofrio et al., 2002).

BoHV-4 has a B-type genome structure, according to ICTV Virus Taxonomy classification of herpesviruses genome (Purdy et al., 2022), characterized by a long unique region (LUR) flanked by polyrepetitive DNA (prDNA) elements, as for Kaposi Sarcoma Associate Herpesvirus (KSKV), Saimirine gammaherpesvirus 2 (SaHV-2), Ateline gammaherpesvirus 3 (AtHV-3), Otarine gammaherpesvirus 4 (OtHV-4) and the others *Gammaherpesvirinae* (McGeoch et al., 2000; Li et al., 2005). Two strains of BoHV-4 have been fully sequenced and annotated so far, the 66-p-347 American strain (Zimmermann et al., 2001) and the V.test European strain (Palmeira et al., 2011), showing a nucleotide identity of 99,55% in average. BoHV-4 genome coding capacity is represented by 79 *orf*s; 59 *orf*s coding for proteins containing conserved domain (CCD) and 20 *orf*s coding for proteins containing non-conserved domain (NCD) (Zimmermann et al., 2001; Palmeira et al., 2011). Using complementary proteomic approaches, a BoHV-4 proteogenomic map, integrating genomic sequencing and proteomic analysis data, was generated by Lété et al; (Lete et al., 2012). The results obtained showed that all 37 identified proteins mapped into previously annotated *orf*s. However, most of these genes and their products have not been structurally and functionally characterized for BoHV-4, among them *orf*45 gene and its product, ORF45 protein. ORF45 is conserved in the *Gammaherpesvirinae* subfamily, there is no *alpha*-, *betaherpesvirinae* and cellular homolog for ORF45, and was characterized only for KHSV (Atyeo and Papp, 2022), MHV-68 (Jia et al., 2005) and RRV (Jia et al., 2005). In the present work, taking advantage of BoHV-4 genome cloned as a Bacterial Artificial Chromosome (BAC) (Donofrio et al., 2008), which allowed us to generate an *orf*45 BoHV-4 null mutant and its pararevertant, BoHV-4 ORF45 was characterized and defined to be *bona fides* ORF45, similarly to KSHV, MHV-68 and RRV ORF45.

## Materials and Methods

### Bovine ORF45 structure prediction

The prediction of the complete ORF45 bovine protein tertiary structure by different ab-initio prediction systems only leaded to low-score confidence structures. However, the N-terminal end of the bovine ORF45 was successfully predicted with the Swiss Model 3D structure prediction server (Waterhouse et al., 2018) by using as a template the homologous portion of the ORF45 protein present in human herpesvirus 8 (PDB: 7opo). The predicted ORF45 bovine structure is 47 residues long (from aminoacid 26 to 72) and displayed a RMSD value of 0.105 A. The Modeller comparative modeling program (Webb and Sali, 2016) has been used to model the bovine ORF45 FxFP motif to the human corresponding one (PDB: 7opm) bound to ERK2. ChimeraX Software was used to display protein structures (Pettersen et al., 2021).

### Cells

HEK (Human Embryo Kidney cells) 293 T (ATCC: CRL-11268), BEK (Bovine Embryo Kidney) (Istituto Zooprofilattico Sperimentale, Brescia, Italy; BS CL-94), BEK *cre*, expressing *cre* recombinase (Donofrio et al., 2008) and Madin–Darby Bovine Kidney (MDBK) cells (ATCC: CRL 6071) were maintained in complete Eagle’s minimal essential medium (cEMEM: 1 mM sodium pyruvate, 2 mM of L-glutamine, 100 IU/ml of penicillin, 100 μg/ml of streptomycin, and 0.25 μg/ml of amphotericin B), supplemented with 10% FBS, and incubated at 37°C/5% CO_2_ in a humidified incubator. All the supplements for the culture medium were purchased from Gibco.

### Constructs Generation

pCMV-ORF45HA was generated amplifying by PCR BoHV-4 ORF45 from pBAC-BoHV-4-A DNA cut with EcoRI. The PCR amplification reaction was carried out in a final volume of 50 μL, containing 20 mM Tris–hydrochloride pH 8.8, 2 mM MgSO4, 10 mM KCl, 10 mM (NH4)2SO4,0.1 mg/mL BSA, 0.1% (v/v) Triton X-100, 5% dimethyl sulfoxide (DMSO), 0.2 mM deoxynucleotide triphosphates, and 0.25 μM of each primer. As a couple of primers were used ORF45 NheI sense and ORF45HA SmaI antisense (see **Supplementary table 1**), to provide BoHV-4-ORF45 with a NheI site at its amino-terminal and a SmaI site and an HA tag at its carboxyterminal. 100 ng of DNA was amplified over 35 cycles, as follows: 1 minute denaturation at 94 °C, 1 minute annealing at 60 °C, and 50 seconds elongation at 72°C with 1U of Pfu recombinant DNA polymerase (Thermo Fisher Scientific). The amplicon ORF45HA was cut with NheI/SmaI and inserted in pEGFP-C1 (Clontech), cut with the same enzyme, to generate pCMV-ORF45HA.

pCMV-ORF45HA was then used as a template to amplify again ORF45HA, with the same PCR parameters described above, with these couple of primers: Fusion-XhoI sense and SmaI-HA-antisense (**Supplementary table 1**). This new amplicon was fused in frame with the GFP ORF, subcloning ORF45, cut with XhoI/SmaI in pEGFP-C1, cut with the same enzymes, generating pEGFP-ORF45-HA.

pTZ-KanaGalK, was generated by sub-cloning the 2232 bp galactokinase prokaryotic expression cassette (GalK), along with the kanamycin resistance expression cassette (Kana), into the pTZ57R shuttle vector (Thermoscientific), cut with KpnI/PstI (Franceschi et al., 2013). The targeting vector, pORF45Left-KanaGalK-RightORF45, was generated firstly by the insertion of the ∼700 bp left ORF45 homology region amplicon (ORF45A sense and antisense, see **Supplementary table 1**) cut with EcoRI/KpnI, in pTZ-KanaGalK, cut with the same enzymes; in this intermediate construct, cut with PstI/HindIII, was subsequently sub-cloned the ∼700 bp right ORF45 homology region amplicon (ORF45B sense and antisense; see **Supplementary table 1**), cut with the same enzymes.

The retargeting vector, pORF45Left-CMVORF45HA-RightORF45 was obtained subcloning the CMVORF45HA-pA entire expression cassette, excised from AseI/MluI cut pCMV-ORF45HA, in pORF45Left-KanaGalK-RightORF45, deprived of KanaGalK selector cassette, through NdeI/MluI restriction digestion.

### Transient Transfection

HEK 293 T cells were seeded into 25cm^2^ flasks (1 × 10^6^ cells/flask) and incubated at 37°C with 5% CO_2_. When cells were sub-confluent, the culture medium was removed and the cells transfected with pCMV-ORF45HA or pEGFP-C1 (Mock control) using Polyethylenimine (PEI) transfection reagent (Polysciences, Inc.). Briefly, 7,5 μg of DNA were mixed with 18,75 μg of PEI (1 mg/ml) (ratio 1:2.5 DNA:PEI) in 500 μl of Dulbecco’s modified essential medium (DMEM) high glucose (Euroclone) without serum. After 15 minutes incubation at room temperature, 2000 μl of medium without serum were added, and the transfection solution was transferred to the cells (monolayer) and left for 6 hours at 37°C with 5% CO2, in a humidified incubator. The transfection mixture was then replaced with fresh cEMEM medium, with 10% FBS, and incubated for 24 hours at 37°C with 5% CO2. To analyse the subcellular localization of BoHV-4 ORF45, HEK 293 T cells were also transfected with pEGFP-ORF45-HA or pEGFP-C1, as a mock control. Twenty four hours after the transfection, the cells were counterstained with DAPI (Thermo Scientific) and observed with a confocal microscope (Leica Microsystems).

### Immunoblotting

Western immunoblotting analysis was performed on protein cell extracts from 25 cm^2^ flasks of HEK 293 T cells transfected with pCMV-ORF45HA or mock transfected. For protein extraction, 100 μl of cell extraction buffer (50 mM Tris–HCl, 150 mM NaCl, and 1% NP-40; pH 8) was added on each pellet and total protein quantification was performed using BCA Protein Assay kit (Pierce™, Thermo Fisher Scientific). Before immunobloting analysis, pCMV-ORF45HA transfected cells extract was enriched in phosphorylated protein, passing through a Phosphoprotein Chelating metal resin, enrichment Kit (Pierce, Thermoscientific), following the protocol suggested by the manufacturers. Different amount of protein samples were electrophoresed on 10% SDS-PAGE and then transferred to PVDF membranes (Millipore, Merck) by electroblotting. The membrane was blocked in 5% skim milk (BD), incubated 1 hour with primary mouse monoclonal antibody anti-HA tag (G036, Abm Inc.) diluted 1:10,000 and then probed with horseradish peroxidase-labeled anti-mouse immunoglobulin (A9044, Sigma), diluted 1:15,000, and finally visualized by enhanced chemiluminescence (Clarity Max Western ECL substrate, Bio-Rad).

### BAC Recombineering and Selection

Recombineering was performed as previously described (Warming et al., 2005) with some modifications. For heat-inducible homologous recombination in SW102 Escherichia coli (*E. coli*), containing BoHV-4-A genome, cloned as a BAC, pBAC-BoHV-4-A, was used the double selector targeting cassette Left-KanaGalK-Right, which was excised from the plasmid backbone pORF45Left-KanaGalK-RightORF45, cut with EcoRI/HindIII. After recombineering, only those colonies that were kanamycin and chloramphenicol positive were kept and grown overnight in 5 ml of LB containing 12.5 μg/ml of chloramphenicol or 50 μg/ml kanamycin. BAC-DNA was purified and analyzed through HindIII restriction enzyme digestion. DNA was then separated by electrophoresis in a 1% agarose gel, stained with ethidium bromide, and visualized through UV light. SW102 bacteria containing BAC-BoHV-4-A-ΔORF45KanaGalK genome were also grown, heat induced and electroporated with HindIII linearized pORF45Left-CMVORF45HA-RightORF45 in the retargeting step. For the counter selection step only the colonies growing on chloramphenicol and not on kanamycin were kept and grown overnight in 5 ml of LB containing 12.5 μg/ml of chloramphenicol. BAC-BoHV-4-A-revORF45HA DNA was purified and analyzed through HindIII restriction enzyme digestion for CMVORF45HA locus fragment targeted integration. Original more detailed protocols for recombineering can also be found at the recombineering website (https://redrecombineering.ncifcrf.gov/).

### Cell Culture Electroporation and Recombinant Virus Reconstitution

BEK or BEK *cre* cells were maintained as a monolayer with cEMEM growth medium with 10% FBS. When cells were sub-confluent (70–90%) they were split to a fresh culture flask (i.e., every 3–5 days) and were incubated at 37°C in a humidified atmosphere of 95% air, 5% CO_2_. pBAC-BoHV-4-A, pBAC-BoHV-4-A-revORF45HA and pBAC-BoHV-4-A-ΔORF45KanaGalK DNAs (5 μg) were electroporated in 600 μl DMEM high without serum (Biorad, Gene Pulser XCell, 270 V, 960 mF, 4-mm gap cuvettes) into BEK and BEK *cre* cells from a confluent 25-cm^2^ flask. Electroporated cells were transferred to new flasks, after 24 hours the medium was replaced with fresh cEMEM, and cells were split 1:2 when they reached confluence at 2 days post-electroporation. Cells were grown until the appearance of cytopathic effect (CPE).

### Viruses and Viral Replication

BoHV-4-A and BAC-BoHV-4-A-revORF45 were propagated by infecting confluent monolayers of BEK or MDBK cells at a multiplicity of infection (M.O.I.) of 0.5 tissue culture infectious doses 50 (TCID_50_) per cell and maintained in cEMEM with only 2% FBS for 2 hours. The medium was then removed and replaced with fresh cEMEM with 10% FBS. When CPE affected the majority of the cell monolayer (∼72 hours post infection), the virus was prepared by freezing and thawing cells three times and pelleting the virions through a 30% sucrose cushion, as previously described (Donofrio et al., 2006). Virus pellets were then resuspended in cold cEMEM without FBS. Viral pellets that were loaded in SDS-PAGE gel were resuspended in 100 μl of cell extraction buffer (50 mM Tris–HCl, 150 mM NaCl, and 1% NP-40; pH 8) and heat denatured. TCID_50_ were determined on BEK cells by limiting dilution.

### Viral Growth Curves

BEK cells were infected with BoHV-4-A and BAC-BoHV-4-A-revORF45 at a M.O.I. of 0.1 TCID_50_/cell and incubated at 37°C for 3 hours. Infected cells were washed with serum-free EMEM and then overlaid with cEMEM with 10% FBS. The supernatants of infected cultures were harvested at scheduled time points (24, 48, 72, and 96 hours post infection), and the amount of infectious virus was determined by limiting dilution on BEK or MDBK cells. Viral titer differences between each time point are the averages of triplicate measurements ± standard errors of the mean (p > 0.05 for all time points as measured by Student’s t-test).

### RNA isolation

Five-millions of cells were resuspended in 1ml of Trizol and stored at -80°C until extraction. Total RNA was isolated by NucleoSpin miRNA kit (Macherey–Nagel, Germany), using the protocol combined with TRIzol lysis (Invitrogen, Carlsbad, CA, USA) and small and large RNA recovery in one fraction. Concentration and quality of RNA were determined by Agilent 2100 (Santa Clara, CA, USA). The isolated RNAs were stored at − 80 °C until use.

### Library preparation and sequencing

RNA samples (RIN >7.5) from four replicates (n = 4) for each condition (Control and Treated) were used for library preparation. RNA-Seq libraries were obtained with the Illumina TruSeq RNA Sample Preparation v2 Kit. Concentration and quality check of libraries were determined by Agilent 2100 Bioanalyzer. Sequencing was performed on a single lane of Illumina HiSeq X (San Diego, CA, USA), 150 cycles paired end.

### Data analysis

RNA-Seq analysis was run with the nf-core/rnaseq v.3.8.1 pipeline (https://nf-co.re/rnaseq). The pipeline integrates TrimGalore v0.6.7 for sequence trimming and STAR v2.7.10a (Dobin et al., 2013) for sequence alignment. Sequences were aligned to the human GRCh38.p13 reference genome. Salmon v1.5.2 (https://combine-lab.github.io/salmon/) was used to quantify alignments to gene regions. The EdgeR Bioconductor package v3.6 (Bioconductor, https://bioconductor.org/packages/release/bioc/html/edgeR.html) was used to estimate differential expression between control and treated samples. Hierarchical cluster analysis was performed with Genesis,(Sturn et al., 2002). Differentially expressed genes (DEGs) were submitted to GO analysis using the Cytoscape (version.3.2.1) plug-in ClueGO (version 2.3.5) (Bindea et al., 2009).

## Results

### BoHV-4 ORF45 has a relatively low protein homology with other *Rhadinovirus* ORF45s and is structurally related to KHSV ORF45

BoHV-4 ORF45 gene, in the genome of BoHV-4, has the same locus position (*orf44-orf45-orf46-orf47*) of the ORF45 gene belonging to the viruses of the same genus, *Rhadinovirus*. However, ORF45 nucleotide sequences and their protein products have often a weak percentage of identity between them. In fact, at the protein level, BoHV-4 ORF45 identity percentage was identified to be 24.9, 21.74, and 23.38 with KSHV, RRV, and MHV68 respectively **(Fig.1)**. ORF45s gene have been sequenced, annotated and their protein products deduced in all *Rhadinovirus* isolated to date, however, only KSHV ORF45 has been characterized in terms of structure (Alexa et al., 2022). BoHV-4 ORF45 protein sequence BLAST analysis identified partial low-score homology sequences and no hits in pdb database. In contrast, HHpred analysis found in KHSV ORF45 N-terminal fragment, containing the binding domain for p90 ribosomal S6 kinase (RSK) and signal-regulated kinase (ERK) complex (Alexa et al., 2022), a significant homology (∼30%) with BoHV-4 ORF45, who was used as a potential structural template **(Fig. 2A)**. Thus, the N-terminal structure of BoHV-4 ORF45 was successfully predicted with Swiss Model 3D prediction server (Waterhouse et al., 2018). As shown in **Fig. 2B**, the predicted BoHV-4 ORF45 structure is well superimposed (RMSD 0.1 A) to the corresponding KHSV ORF45 orthologue bound to RSK2 kinase **(Fig. 2B)**. Further, the key Val and Phe (VP) interacting motif residues are also conserved in the BoHV-4 ORF45 and well fits the hydrophobic pocket located in the N-terminal of AGC-type kinase domain (NTK) of RSK **(Fig. 2C)**. Therefore, BoHV-4 ORF45 protein is expected to bind to phosphorylated ERK2, since the substrate-mimicking FxFP motif present in the KHSV ORF45 is also conserved in the BoHV-4 ORF45 and well fits the hydrophobic F-site of ERK2 **(Fig. 2D)**.

**Fig. 1.**
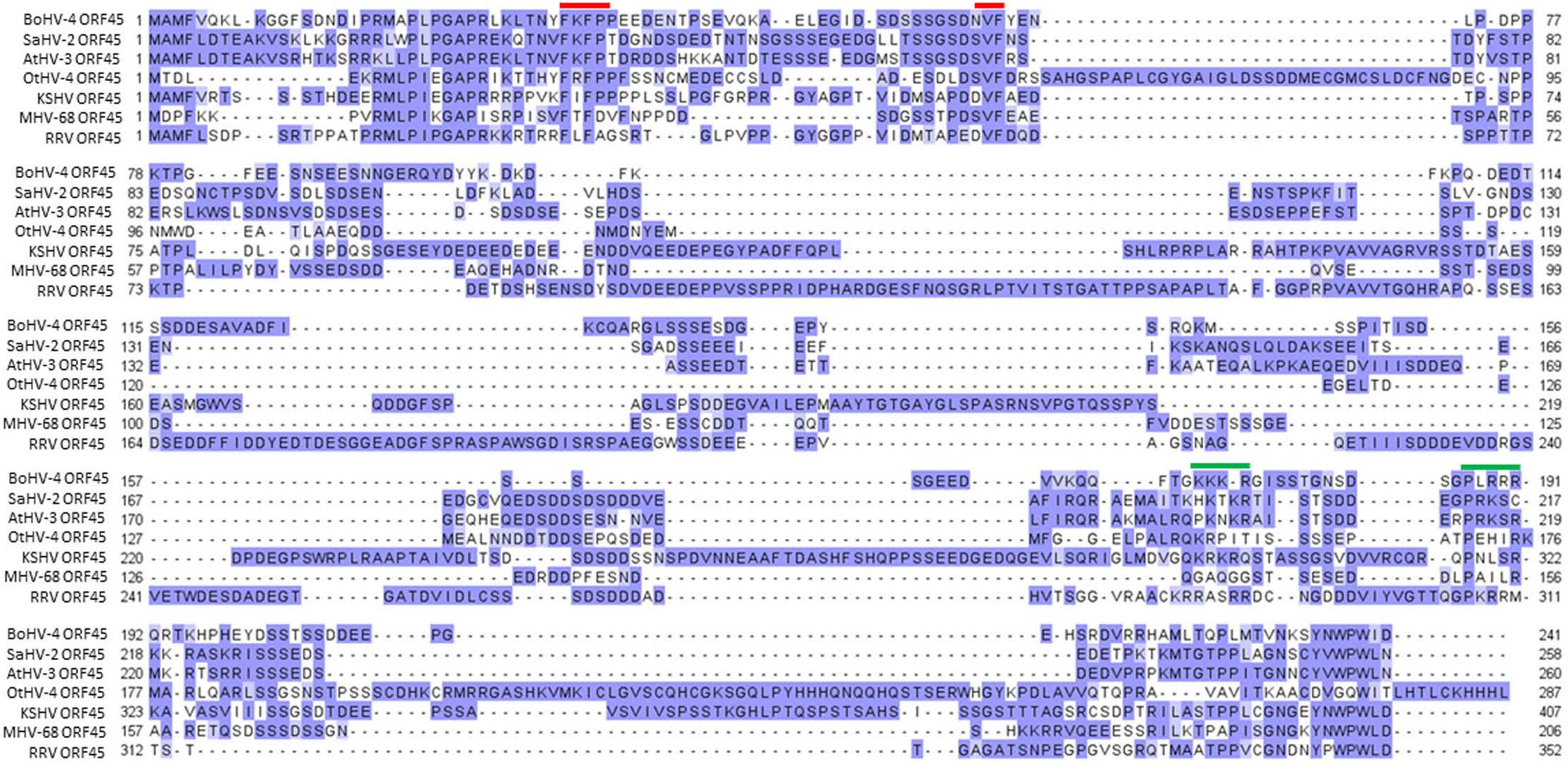
Alignment of representative *Rhadinovirus* ORF45 protein sequesnces. Alignment of the bovine gammaherpesvirus 4 ORF45 protein (BoHV-4 ORF45; Acc. n. AEL29789.1) sequence with a subset of protein homologs, hypothetical protein Saimirine gammaherpesvirus 2 (SaHV-2 ORF45; Acc. n. CAC84340.1), Ateline gammaherpesvirus 3 (AtHV-3 orf45; Acc. n. NP_048018.1), Otarine gammaherpesvirus 4 (OtHV-4 ORF45; Acc. n. QRE02526.1), Kaposi Sarcoma Associate Herpesvirus (KSHV ORF45; Acc. n. BAV17895.1), Murid herpesvirus 4 (MHV-68 ORF45; NP_044882.1), Rhesus monkey rhadinovirus (RRV ORF45; Acc. n. AAF60024.1). Conserved amino acids are drawn according to their percentage similarity with the consensus sequence (>80% dark-blue, >60% medium blue, >40% light-blue, <40% white). Conserved functional sites and nuclear localization signals are marked with red and green bars, respectively.

**Fig. 2.**
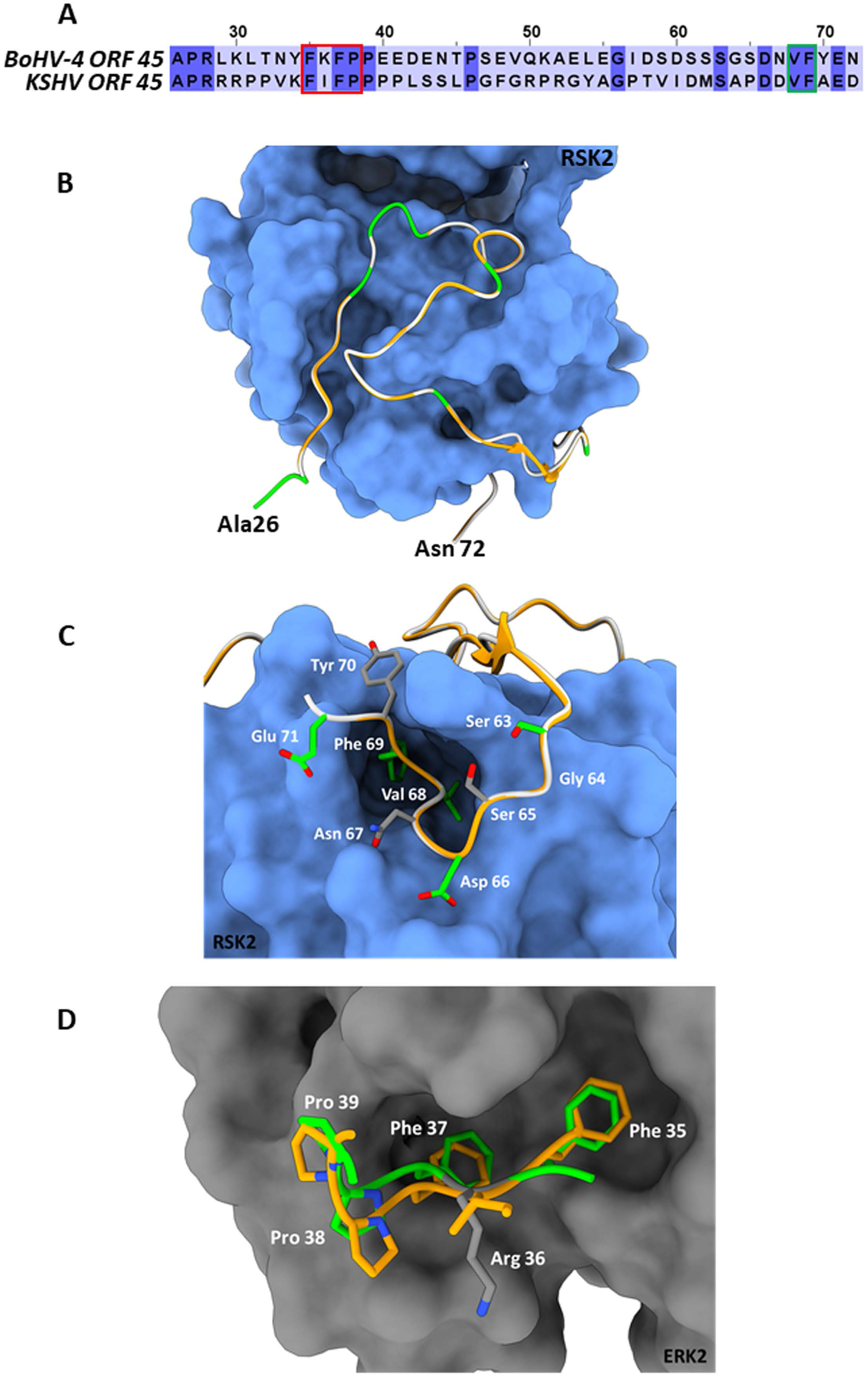
BoHV-4 ORF45 predicted structure. **A**) Amino acid sequence alignment between the BoHV-4 ORF45 (upper sequence, from aa. 26 to 72) and the corresponding residues of KHSV ORF45 hortologue (lower sequence). The conserved amino acids are shown on a dark-blue background. The red and the green rectangles indicate the ORF45 VP-interacting motif and the substrate-mimicking FxFP motif, respectively. **B**) The predicted Ca backbone of BoHV-4 ORF45 (in green the conserved residues; in white the non-conserved residues) is superimposed onto the human ORF45 Ca structure (in orange) bound to RSK2 (protein surface in blue) (PDB: 7opm). The N- and C-termini of the bovine protein are labelled. **C**) Particular of the predicted BoHV-4 ORF45 3D structure (Ca and side chains) superimposed onto Ca KHSV ORF45 Val-Phe binding motif bound to RSK2. Color code of Ca chains as in “B”, while the bovine ORF45 side chains are rendered in in green/CPK or gray/CPK sticks, if conserved or non-conserved, respectively. **D**) Superposition of the predicted BoHV-4 ORF45 3D structure FxFP motif onto the corresponding human ORF45 (PDB: 7opm) residues bound to the hydrophobic F-site of ERK2 (protein surface in grey) (PDB: 7opm). Human ORF45 Ca and side chains are in orange/CPK sticks, the bovine Ca and side chain residues are in green/CPK or gray/CPK sticks if conserved or non-conserved, respectively.

### BoHV-4 ORF45 is a phosphoprotein

BoHV-4 ORF45 has an *in silico* predicted length, starting from its nucleotide sequence (*orf45)* (Zimmermann et al., 2001; Palmeira et al., 2011), of 241 aminoacidic residues, an isoelectric point (IP) of 4.76, a mass of 27.142 kDa, an aliphatic index of 40.46 and is lacking a putative signal peptide. Hence, in general, BoHV-4 ORF45 could be described as an intracellular soluble acidic protein. Since BoHV-4 ORF45 has not been characterized so far and consequently no specific antibodies have been developed, to follow its expression in mammalian cells, a carboxyterminal HA tagged ORF45, pCMV-ORF45HA, was constructed. Surprisingly, BoHV-4 ORF45 had an SDS-PAGE migration almost double than predicted, ∼55 kDa **(Fig. 3A)**. This is probably due to some post-transcriptional modifications and in agreement with what was found for MHV68 ORF45 (Jia et al., 2005) and KSHV ORF45 (Zhu and Yuan, 2003). Glycosylation sites were not predicted and not identified when BoHV-4 ORF45 was treated with endoglycosidases (data not shown). However, NetPhos-3.1 (Blom et al., 1999; Blom et al., 2004) BoHV-4 ORF45 analysis indicated the presence of several phosphorylation sites, where serine and threonine were mainly involved, anyhow, although with a lower frequency and score, also tyrosine **(Fig. 3B)**. Finally, BoHV-4 ORF45 phosphorylation, in pCMV-ORF45HA transfected cells extracts, was confirmed by a phosphoprotein affinity resin which was able to retain BoHV-4 ORF45 **(Fig. 3 C)**.

**Fig. 3.**
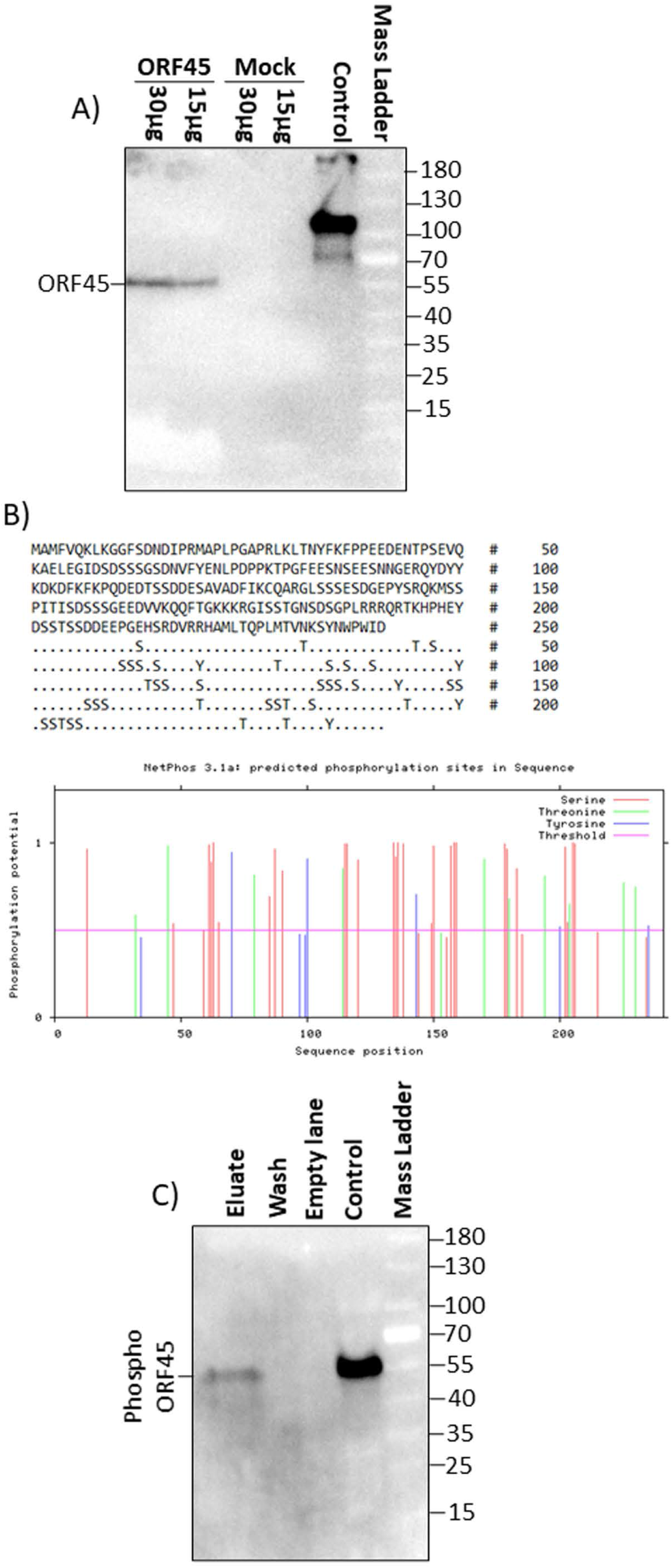
BoHV-4 ORF 45 phosphorylation. **A)** Western immunoblotting of pCMV-ORF45HA (ORF45) and pEGFP-C1 (Mock) transfected cells protein extracts (30 and 15 μg of total protein extract). A positive antibody control was established with an HA tagged unrelated protein. **B)** BoHV-4 ORF45 protein sequence along with Serine (S) Threonine (T) and Tyrosine (Y) residues potentially phosphorylated as predicted by NethPhos 3.1, where the scores above 0.500 indicates positive predictions. **C)** Western immunoblotting of concentrated eluate and wash of pCMV-ORF45HA transfected cells extract, passed through a Phosphoprotein Chelating metal resin. A positive antibody control was established with pCMV-ORF45HA transfected cells extract.

### BoHV-4 ORF45 localizes to the cell nucleus

As predicted by PSORT sequence analysis (Nakai, 2000) a nuclear localization signal was identified in BoHV-4 ORF45 (78%; K=9/23; consensus: KKKR at 172 and PLRRRQR at 187) but not a nuclear export signal as detected in KSHV ORF45 (Li and Zhu, 2009). For examining the capability of BoHV-4 ORF45 to localize to the nucleus, independently from other viral protein backgrounds, GFP *orf* was fused in frame with BoHV-4 HA tagged ORF45 *orf* and a plasmid construct, pEGFP-ORF45-HA, was generated. Twenty-four hours pEGFP-ORF45-HA post-transfected cells were visualized with a confocal microscope, as predicted, GFP fluorescent signal was well observed within the cell nuclei **(Fig. 4A, B, C and D)**. Therefore, BoHV-4 ORF45 was able to deliver GFP, known to be a cytoplasmic protein (**Fig. 4D)**, into the cell nucleus.

**Fig. 4.**
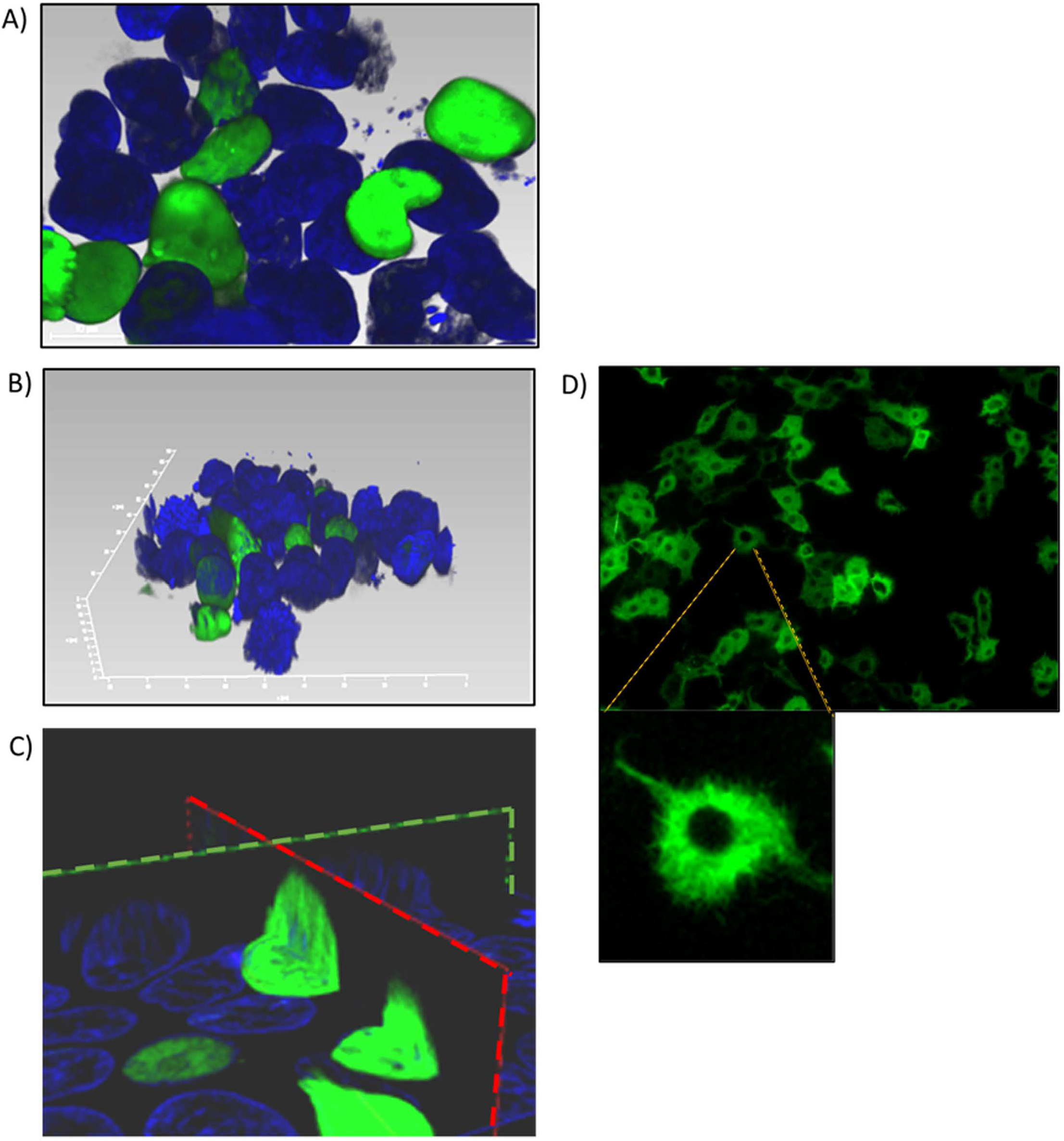
Subcellular localization of BoHV-4 ORF45. Cells were transfected with a construct, pEGFP-ORF45-HA, where GFP *orf* was fused to the 5’ end of BoHV-4ORF45 *orf*. Transfected cells were counterstained with DAPI and observed with a confocal microscope at 24 hours post transfection. **A)** 2D image of pEGFP-ORF45-HA transfected cells where GFP-ORF45 is localized within the nuclei (green). **B)** 3D image of cell nuclei containing GFP-ORF45 (green). **C)** Sagittal (green dashed line) and transverse (red dashed line) image of cell nuclei containing GFP-ORF45. **D)** 2D image of pEGFP-C1 transfected cells control, where GFP is only localized within the cytoplasm (green).

### BoHV-4 ORF 45 is an essential gene for BoHV-4 lytic replication and is associated with the virus

To investigate the role of the BoHV-4 ORF45 during viral lytic replication, the BoHV-4 ORF45 gene was deleted by site-specific insertional mutagenesis mediated by heat inducible homologous recombination (Warming et al., 2005) in the genome of a BoHV-4 cloned as a BAC (BAC-BoHV-4-A). BAC-BoHV-4-A was originally derived from a non-pathogenic strain of BoHV-4 isolated from the milk cellular fraction of a healthy cow (Donofrio et al., 2008). The targeting cassette, Left-KanaGalK-Right, containing the 2232-bp KanaGalK DNA stuffer double selecting cassette (Donofrio et al., 2007) flanked by two BoHV-4 ORF45 gene homologous flanking regions, was generated to mediate insertion and deletion of the BoHV-4 ORF45 coding region. BoHV-4 ORF45 gene is transcribed from the opposite BoHV-4 genome DNA strand (Zimmermann et al., 2001; Palmeira et al., 2011) like the ORF46 but in contrast to ORF44 **(Fig. 5A)**. Therefore, although a large deletion and insertion were made, this should not affect ORF44 and ORF46 transcription/translation and the viral phenotype obtained should be exclusively due to the loss of ORF45. The Left-KanaGalK-Right targeting cassette was excised from the plasmid back-bone and electroporated into SW102 *E. coli* containing pBAC-BoHV-4-A, to generate pBAC-BoHV-4-A-ΔORF45KanaGalK **(Fig. 5B)**. Selected targeted clones were analyzed by PCR, sequencing (data not shown) and HindIII restriction enzyme digestion **(Fig. 5D)**. Further, to stress that the authenticity of pBAC-BoHV-4-A-KΔORF45KanaGal phenotype was exclusively due to the loss of ORF45, and not a mere artifact, a control BoHV-4 with a carboxyterminal HA tagged ORF 45, transcriptionally driven in an opposite direction, respect to the natural ORF45, by a heterologous promoter (CMV), was generated. The retargeting cassette, Left-CMV-ORF45HA-Right, was excised out from the plasmid backbone and electroporated into SW102 E. *coli* containing pBAC-BoHV-4-A-ΔORF45KanaGalK and pBAC-BoHV-4-A-revORF45HA was obtained **(Fig. 5C)**. Again, selected targeted clones were analyzed by PCR, sequencing (data not shown) and HindIII restriction enzyme digestion **(Fig. 5E)**. When pBAC-BoHV-4-A-ΔORF45KanaGalK and pBAC-BoHV-4-A-revORF45HA were electroporated into BEK or BEK/cre cells, to excise out the BAC cassette, plaques from the viable virus were obtained only on pBAC-BoHV-4-A-revORF45HA transfected cells monolayers but not on those transfected with pBAC-BoHV-4-A-ΔORF45KanaGalK **(Figure 6A)**. ORF45 deletion rendered BoHV-4-A unable to be reconstituted and productively replicated, such phenotype was rescued by CMV-ORF45HA expression cassette **(Fig. 6B)**, thus showing ORF45 indispensability in the contest of BoHV-4 lytic replication and in line with data obtained for KHSV ORF45 (Fu et al., 2015) and MHV68 OFR45 (Jia et al., 2005). Since BoHV-4-A-revORF45HA infectious virus produced an HA-tagged form of ORF45, detectable by an anti-HA mAb **(Fig. 6C)**, it was of interest to know if BoHV-4 ORF45 is associated with the virus as previously observed by Palmeira et al., (Lete et al., 2012) in a proteomic analysis setting. As expected, when CsCl_2_ gradient purified virus was analyzed by SDS-PAGE and western blotting, with an anti HA mAb, a specific band corresponding to ORF 45, was well evidenced **(Fig. 6D)**.

**Fig. 5.**
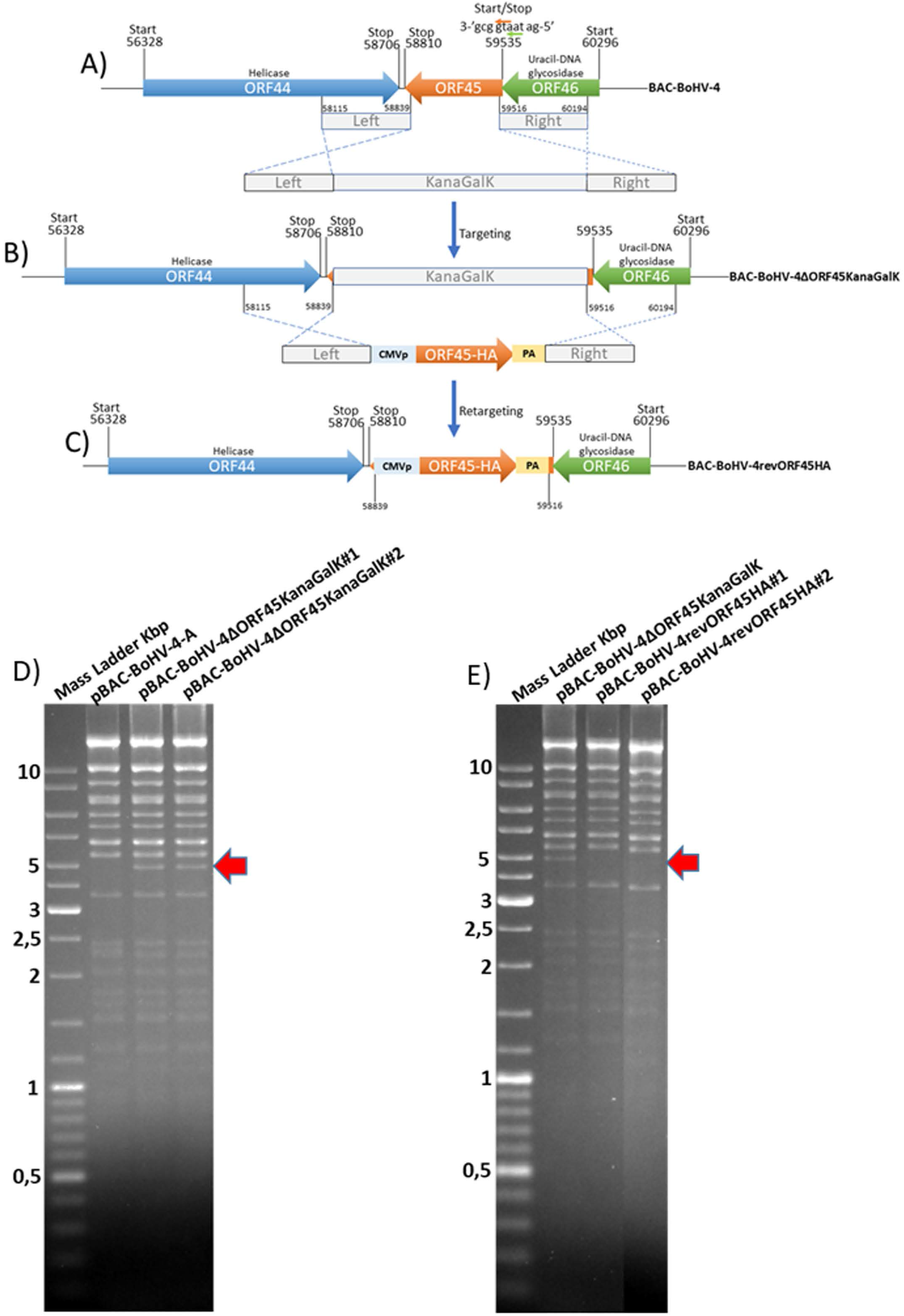
Overall strategy to disrupt BoHV-4 ORF45 gene via heat inducible homologous recombination. **(A and B)** Diagram (not to scale) of the BoHV-4 ORF45 gene (orange arrow; nucleotide 59535 and 58810) positioned between ORF 44 (blue arrow; nucleotide 56328 and 58706) and ORF46 (green arrow; nucleotide 60296 and 59535). The last nucleotide of the ORF46 stop codon and the first nucleotide of the ORF45 start codon (a) is overlapped. ORF46 and ORF45 are transcribed on the opposite direction respect to the ORF44 [based on the complete genome published sequence (GenBank accession number AF318573)]. The 2232-bp Kana/GalK selectable DNA stuffer (gray), flanked by left (nucleotide 58115 and 58839; 724 bp) and right homologous regions (nucleotide 58516 and 60194; 678 bp), was introduced between the positions 58839 and 59516, deleting most of the ORF45 sequence but leaving intact ORF44 and ORF46 and BoHV-4-A-ΔORF45KanaGalK was generated. **B)** CMVORF45HA expression cassette flanked by the left and right homologous regions (grey) was used to replace 2232-bp Kana/GalK selectable DNA stuffer (gray) and BoHV-4-A-revORF45HA was so generated **C). D)** HindIII restriction enzyme profile of two representative pBAC-BoHV-4-A-ΔORF45KanaGalK clones (#1 and #2) compared with the parental pBAC-BoHV-4-A. The diagnostic band, indicated by a red arrow, was well observable in pBAC-BoHV-4-A-ΔORF45KanaGalK respect to pBAC-BoHV-4-A. **E)** HindIII restriction enzyme profile of two representative pBAC-BoHV-4revORF45HA clones (#1 and #2) compared with the derivative pBAC-BoHV-4-A-ΔORF45KanaGalK. The the missing band, indicated by a red arrow, was well observable in pBAC-BoHV-4-A-revORF45HA respect to pBAC-BoHV-4-A-ΔORF45KanaGalK.

**Fig. 6.**
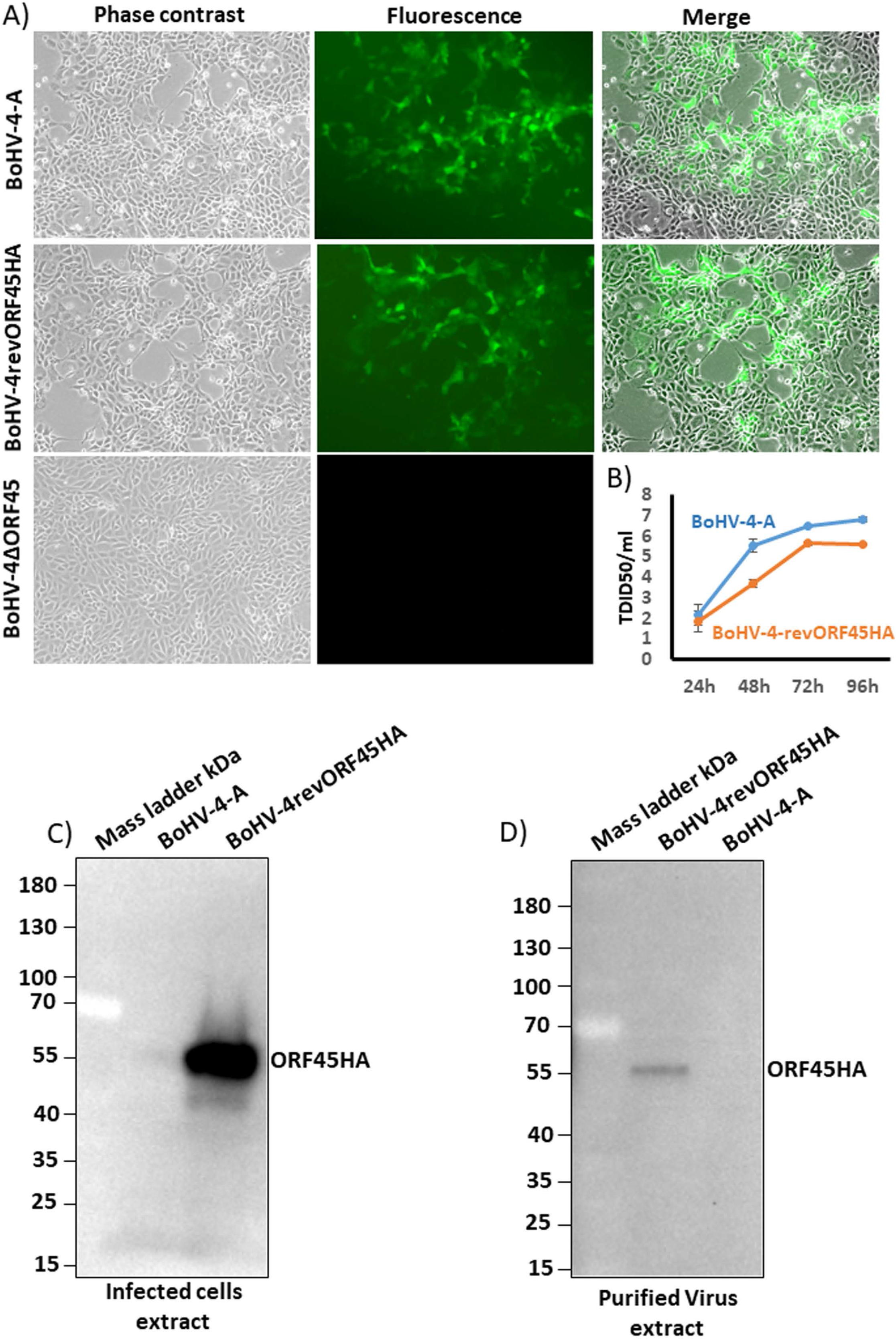
BoHV-4 recombinants reconstitution and replication. **A)** Representative images (phase contrast, fluorescence and merged; 10×) of BEK cells electroporated with pBAC-BoHV-4-A, pBAC-BoHV-4-A-revORF45HA and pBAC-BoHV-4-A-ΔORF45KanaGalK. CPE induced by reconstitution of IRVPs is recognizable only for pBAC-BoHV-4- and pBAC-BoHV-4-A-revORF45HA as revealed by green plaques. **B)** Replication kinetics of BoHV-4-A and BAC-BoHV-4-A-revORF45. **C)** Western immunoblotting of BoHV-4-A and BoHV-4-A-revORF45HA infected cells extract. **D)** Western immunoblotting of BoHV-4-A and BoHV-4-A-revORF45HA purified virus extract. All experiments were repeated three times with identical results.

### BoHV-4 ORF45 expression alters the cellular transcriptome

Since BoHV-4 ORF45 possesses an acidic domain (between amino acids 40 and 66), is phosphorylated, and localizes into the cell nucleus, these features would suggest that BoHV-4 ORF45 could also take part, perhaps indirectly through protein/protein interactions with other transcription factors, in a transcriptional regulatory network. To test this hypothesis, a comparative transcriptome analysis of cells expressing BoHV-4 ORF45 versus BoHV-4 ORF45 un-expressing cells was performed.

RNA-seq analysis generated an average number of 55.8 M (111.56 M paired reads) reads per sample, with about 81.4% of the total reads that were correctly mapped on the human reference genome (**Supplementary table 2**, for statistics). A total of 30738 unique genes were present in both groups, whether 94 and 190 genes were only detected in treated and control samples, respectively **(Fig. 7A)**. The principal component analysis (PCA) clearly separates control vs treated samples **(Fig. 7B)**. A total of 984 differentially expressed genes (DEGs), (FDR < 0.05), were identified **(Fig.7C and Supplementary table 3)**. Biological pathway analysis was performed for high significant DEGs (n=113, FDR < 0.01 and LogFC<|0.5|). Gene Ontology analysis (GO) showed variation in genes involved in mitotic DNA damage checkpoint, response to DNA damage by p53 and pathway restricted to SMAD phosphorylation **(Fig. 8A and Supplementary table 4)**. Reactome pathway database interrogation identified genes related to TP53 regulation of cell cycle, kinase-mediated activation of nuclear transcription and DNA damage senescence response **(Fig. 8B and Supplementary table 4)**. This in complete agreement with KSHV ORF45 functions (Atyeo and Papp, 2022) and interaction with p90 ribosomal S6 kinase (RSK) and signal-regulated kinase (ERK) complex.

**Fig. 7.**
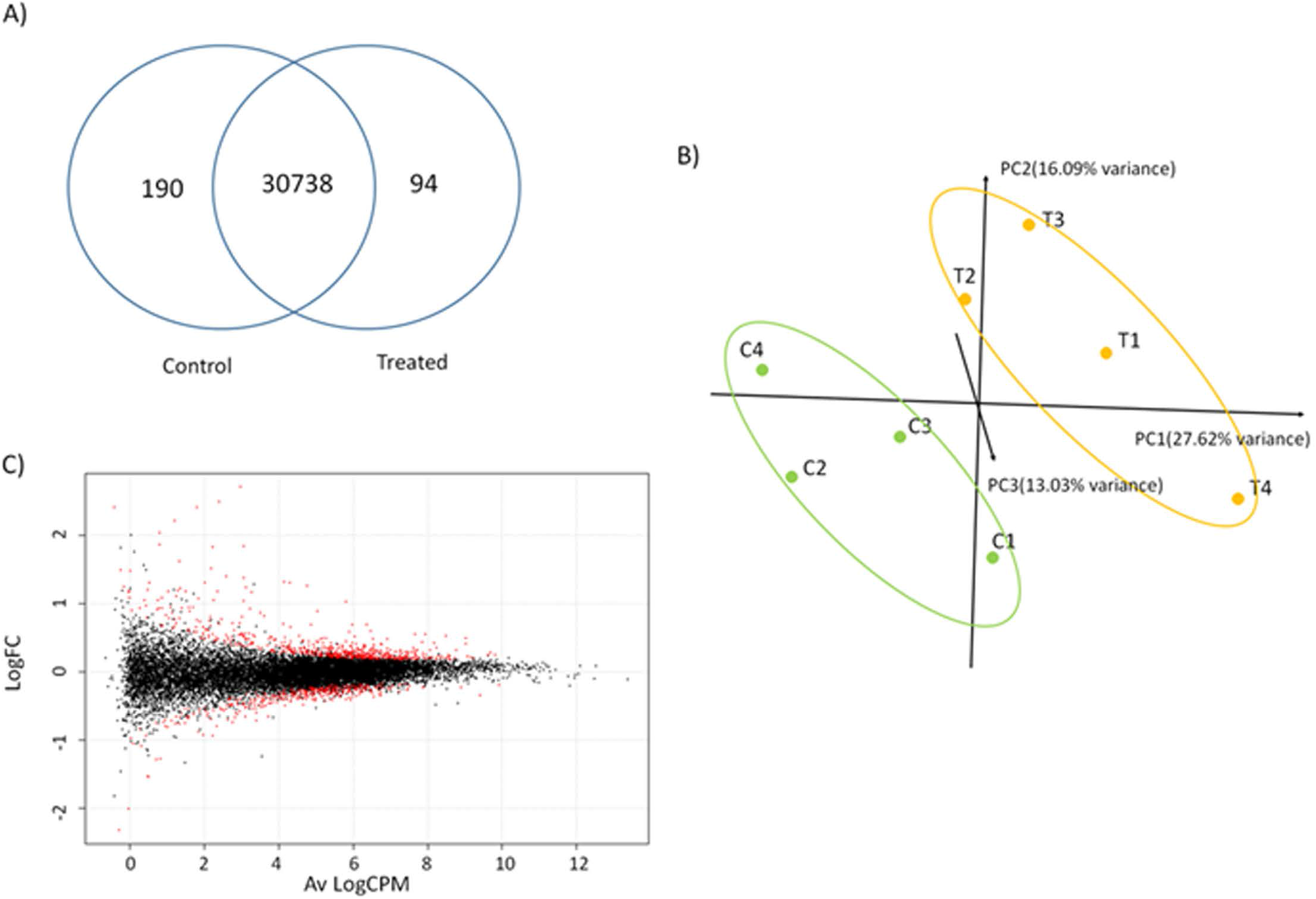
Transcriptome analysis. **A)** Shared and unique genes identified in control and treated samples; **B)** Principal Component Analysis of normalized reads for treated (T1-T4) and control (C1-C4) samples; **C)** Smear-Plot representing the average logarithmic count per millions (Av LogCPM) gene expression and logaritmic fold change variation (LogFC) between control and treated samples calculated with EdgeR. In red Differentially expressed genes (DEGs).

**Fig. 8.**
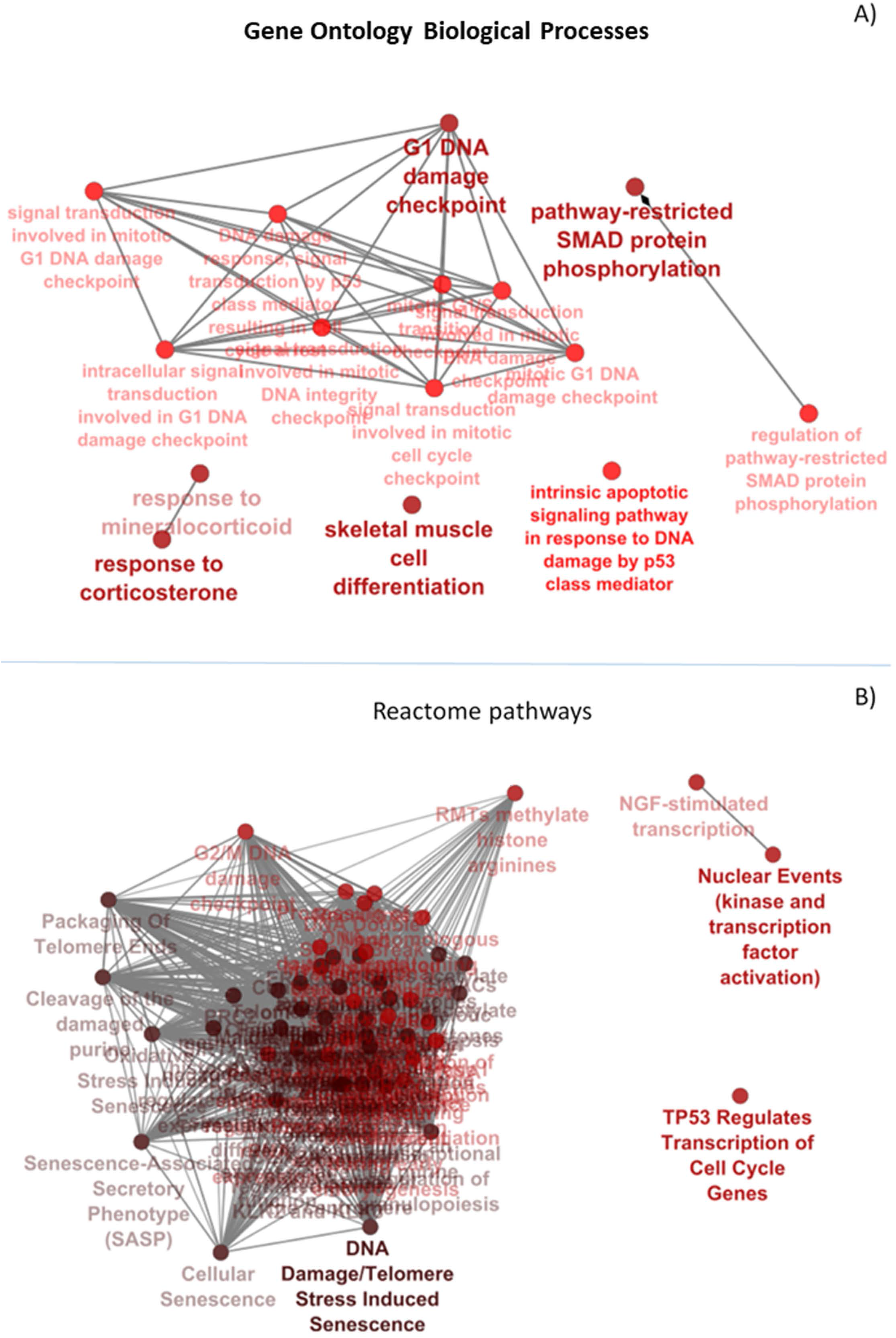
Representation of significant enriched terms found for: A) Biologycal processes in Gene Ontology (GO) analysis and **B)** Reactome pathways for DEGs (FDR<0.01, LogFC|0.5|)

## Discussion

A herpesviral virion is generally composed by 3 electron microscopy morphologically distinct structures: an envelope, a capsid and an electron dense material defined as the tegument, located between the envelope and the capsid (Roizmann et al., 1992). Proteomic analysis of BoHV-4 mature virions, identified 5 envelope proteins, 9 capsid proteins, and 13 tegument proteins (Lete et al., 2012). Among the tegument proteins, ORF45 was one of the most abundant. The criteria utilized to characterize BoHV-4 tegument proteins were based on resistance to protease digestion in absence of detergent and susceptibility to protease digestion following treatment with envelope dissolving detergent. BoHV-4 ORF45 sequence similarity with other previously characterized *Rhadinovirus* ORF45s were not strong enough for attributing with certainty structural and functional characteristics of an authentic ORF45, mainly in absence of experimental data. In this work, when BoHV-4 ORF45 amino terminal region was compared with that of KHSV ORF45, it allowed to predict a structure with potential similar functions and interactive pathways to those of KHSV ORF45. Overexpression of HA tagged or GFP fused BoHV-4 ORF45 was identified as a phosphoprotein who localizes to the host cell nucleus as it was the case for KSHV, RRV and MHV68. By construction and analysis of a BoHV-4 ORF45-null mutant ad its pararevertant, using BAC system, was demonstrated that the ORF45 null mutant was uncapable of virions production, indicating that newly synthetized ORF45 is essential for BoHV-4 lytic replication and such defect could be rescued *in cis* by reintroducing ORF45. Moreover, the HA tagged ORF45 pararevertant virus allowed to experimentally demonstrate that BoHV-4 ORF45 is associated with the virus particle, corroborating the notion, previously partially demonstrated (Lete et al., 2012), of BoHV-4 ORF45 as part of the tegument. Tegument proteins are not just structural viral proteins essentially implicated in viral entry, replication, morphogenesis, and egress. They are the first viral proteins to be delivered inside the host cells immediately after virus attachment, membrane fusion and uncoating. They can specifically modify, for increasing infection fitness, the host intracellular environment during the establishment of *de novo* infection and modify, through a specific interactome, host signaling, epigenetic and transcriptome (Sathish et al., 2012; Atyeo and Papp, 2022). KSHV ORF45 is one of the most, if not the most, investigated *Gammaherpesvirus* tegument protein, as well as its interaction with p90 ribosomal S6 kinase (RSK) and signal-regulated kinase (ERK) complex (RSK/ERK). Since BoHV-4 ORF45 structure was well superimposed (RMSD 0.1 A) to the corresponding KHSV ORF45 orthologue, the key Val and Phe (VP) interacting motif residues are also conserved in the BoHV-4 ORF45 and well fits the hydrophobic pocket located in the N-terminal of AGC-type kinase domain (NTK) of RSK, similar function for BoHV-4 ORF45 could be envisioned. ORF45 sustains activation of the p90 ribosomal S6 kinase (RSK) and signal-regulated kinase (ERK) MAP kinase pathway by binding to the RSK/ERK complex and preventing their dephosphorylation (Kuang et al., 2009; Alexa et al., 2022) or acting as a SUMO ligase and SUMOylates RSK to promote its kinase activity (Liu et al., 2022). One of the downstream targets of phosphorylated RSK/ERK complex is the cellular transcription by direct phosphorylation of transcription factors. RSKs regulate several transcription factors, including CREB, serum response factor (SRF), ER81, estrogen receptor-α (ERα), nuclear factor-κB (NF-κB), NFATc4, NFAT3 and the transcription initiation factor TIF1A (Romeo et al., 2012). Starting from this information, it was of interest to investigate the impact of BoHV-4 ORF45 on cellular transcriptome, something which was little studied or not at all for ORF45 belonging to other *Gammaherpesvirus*. Applying a stringent selection, (DEGs; FDR < 0.01 and LogFC<|0.5|) for biological pathways identification, gene ontology analysis, and reactome pathways database interrogation, a strong transcriptome alteration was identified in cells expressing BoHV-4 ORF45. Among the most remarkable, stress response, cell differentiation, pathways-restricted to SMAD protein phosphorylation, TP53 regulates transcription of cell cycle genes and DNA damage checkpoint pathways, in cells expressing BoHV-4 ORF45 that are consistent with cellular functional modulation in response RSK/ERK activation. Indeed, RSK binds and enhances the function of the transcriptional co-activators CREB-binding protein (CBP) and p300 (Nakajima et al., 1996; Wang et al., 2003). These proteins are large homologous molecules that facilitate complex formation between different components of the transcriptional machinery. Among the transcription factors that associate with CBP and p300 are cAMP response element-binding protein (CREB), Fos Proto-Oncogene, AP-1 Transcription Factor Subunit FOS, Jun Proto-Oncogene, AP-1 Transcription Factor Subunit JUN, Signal Transducer and Activator of Transcription (STAT), Myogenic Differentiation protein (MyoD), E2F Transcription Factor (E2F), NF-κB and steroid receptors including Estrogen receptor alpha (Erα) (Romeo et al., 2012). Among the top 200 most significant DEGs, CREB5, FOS, FOSL1, JUN, NFKBIL1 genes showed variation in two group of C and T samples. These transcription factors could be directly and indirectly associated to one of the pathways transcriptionally impacted by BoHV-4 ORF45. For instance, MyoD with cell differentiation (Noda et al., 2009); CREB/ FOS/JUN with TP53 regulates transcription of cell cycle genes (Schreiber et al., 1999; Giebler et al., 2000); NF-κB and CREB with pathways-restricted to SMAD (Freudlsperger et al., 2013); E2F with cell cycle progression, DNA repair, replication, and G/M checkpoints (Ren et al., 2002); CREB/Erα with corticosteroids stress response (Jing et al., 2016). Finally, we also observed up-regulation of the ETS Variant Transcription Factors, ETV4 and ETV5, in cells expressing BoHV-4 ORF45, likely associated to RSK/ERK activation. A previous study in cancer cell lines reported that αvβ3 integrin signaling may activate ERK pathway, resulting in ETV4 transcription, PD-L1/L2 expression, and evasion attacks from the immune system (Ma et al., 2021). Expression of endogenous retroviruses envelope protein in different human cancer cell lines was observed to induce the transcription of ETV4 and ETV5 which are downstream effectors of the MAPK ERK1/2 (Lemaitre et al., 2017).

In conclusion, in the present paper, for the first time, a general characterization from the structural and functional point of view of BoHV-4 ORF45 was obtained, posing the base for further investigation on each of the characteristics pointed out here. Although in general, BoHV-4 ORF45 characteristics can be ascribed to those of KHSV ORF45.

## Acknowledgments and founding

This research was supported by internal found of Parma University.

## Author contributions

G.D. designed the experiments. L.R., E.C., V.F., R.S., D.C., B.L. and G.D. performed the experiments and analyzed the data. G.D. wrote the manuscript and supervised the project.

