## Supplementary Figure 1 for "Characterization of BoHV-4 ORF45"

**Table 1: List of primers used in this work**

| Primer name | Sequence 5'-3' |
|---|---|
| ORF45 A sense | CCCGAATTCTGGGACATCTTTTCTTCAAAAAAACTTTGT |
| ORF45 A antisense | CCCGGTACCCATATGAAAAGTTACAATTGGCCATGGATTGACTGA |
| ORF45 B sense | CCCCTGCAGACGCGTCCCCTTGAGCTTCTGCACAAACATCGCCAT |
| ORF45 B antisense | CCCAAGCTTCTACAGTTATCCCCATTTATGAAACAAAAA |
| ORF45NheI sense | GGGGCTAGCCCACCATGGCGATGTTTGTGCAGAAG |
| ORF45HA SmaI anti | CCCCCCGGGTTAGGCGTAGTCGGGCACGTCGTAGGGGTAGTCAATCCATGGCCAATTGTAACTTTT |
| Fusion XhoI sense | CTCAGATCTCGAGCTGCGATGTTTGTGCAGAAGCTCAAGGGGGG |
