## Supplementary Figure 2 for "Characterization of BoHV-4 ORF45"

| | Read 1 | Read 2 | %Uniquely mapped | %Mapped to multiple loci | %Mapped to too many loci | %Unmapped: too short | % Unmapped: other |
|---|---|---|---|---|---|---|---|
| C1 | 55196577 | 55196577 | 77.08 | 3.51 | 0.01 | 19.36 | 0.05 |
| C2 | 45835201 | 45835201 | 78.71 | 3.36 | 0.01 | 17.86 | 0.06 |
| C3 | 61841774 | 61841774 | 75.71 | 3.98 | 0.01 | 20.25 | 0.05 |
| C4 | 76628130 | 76628130 | 79.11 | 3.41 | 0.01 | 17.41 | 0.06 |
| T1 | 41395922 | 41395922 | 85.15 | 3.55 | 0.01 | 11.24 | 0.06 |
| T2 | 84994155 | 84994155 | 85.82 | 3.93 | 0.01 | 10.18 | 0.06 |
| T3 | 38229254 | 38229254 | 84.54 | 4.16 | 0.01 | 11.23 | 0.06 |
| T4 | 42048047 | 42048047 | 84.80 | 3.47 | 0.01 | 11.66 | 0.06 |

**Supplementary Table 2**: Sequencing statistics
