## Supplementary Figure 3 for "Characterization of BoHV-4 ORF45"

**Supplementary Table 3.** List of Differentially Expressed Genes (DEGs) between control and treated samples (FDR<0.05). Gene name (gene_name), gene identification code (gene_id), logarithmic value of fold change expression variation between two groups (logFC) and False Discovery Rate value for significance (FDR) were reported.

| gene_name | gene_id | logFC | FDR |
|---|---|---|---|
| H1-2 | ENSG00000187837 | 1,250184 | 3,69E-48 |
| DHRS2 | ENSG00000100867 | 1,306915 | 2,12E-46 |
| ENSG00000289609 | ENSG00000289609 | 1,833162 | 4,63E-46 |
| ETV4 | ENSG00000175832 | 2,39812 | 6,62E-36 |
| ETV5 | ENSG00000244405 | 2,483144 | 3,49E-32 |
| LIF | ENSG00000128342 | 0,878307 | 1,21E-27 |
| CYP4F2 | ENSG00000186115 | 1,815211 | 3,48E-25 |
| BTG2 | ENSG00000159388 | 0,592108 | 4,28E-23 |
| MAFF | ENSG00000185022 | 1,226604 | 7,11E-22 |
| ENSG00000260103 | ENSG00000260103 | 1,510454 | 9,85E-20 |
| ENSG00000288758 | ENSG00000288758 | 2,206582 | 1,21E-19 |
| CYP4F3 | ENSG00000186529 | 1,396256 | 1,32E-19 |
| AEN | ENSG00000181026 | 0,52144 | 1,75E-17 |
| DDB2 | ENSG00000134574 | 0,683171 | 6,60E-17 |
| PHLDA3 | ENSG00000174307 | 0,679982 | 6,60E-17 |
| GDF15 | ENSG00000130513 | 1,033762 | 2,42E-16 |
| CDKN1A | ENSG00000124762 | 0,579161 | 7,89E-16 |
| SPRY4 | ENSG00000187678 | 2,030078 | 7,89E-16 |
| ACTA2 | ENSG00000107796 | 0,679625 | 5,38E-15 |
| SNHG17 | ENSG00000196756 | 0,548122 | 6,55E-15 |
| ENSG00000289688 | ENSG00000289688 | 1,854089 | 2,30E-13 |
| GPR3 | ENSG00000181773 | 1,169428 | 7,56E-13 |
| H1-0 | ENSG00000189060 | 0,488185 | 9,95E-13 |
| ENSG00000287064 | ENSG00000287064 | 1,605897 | 1,03E-12 |
| INPP5D | ENSG00000168918 | 0,808769 | 1,04E-12 |
| SESN2 | ENSG00000130766 | 0,430181 | 8,52E-12 |
| SNHG15 | ENSG00000232956 | 0,539891 | 1,61E-11 |
| ATF3 | ENSG00000162772 | 0,421157 | 2,14E-11 |
| PLK3 | ENSG00000173846 | 0,69972 | 1,37E-10 |
| FOSL1 | ENSG00000175592 | 0,920662 | 1,81E-10 |
| SNHG12 | ENSG00000197989 | 0,478966 | 3,61E-10 |
| ZNF79 | ENSG00000196152 | 0,484098 | 1,13E-09 |
| H2BC11 | ENSG00000124635 | 1,292668 | 1,85E-09 |
| PVT1 | ENSG00000249859 | 0,48707 | 1,94E-09 |
| CREB5 | ENSG00000146592 | 0,471642 | 2,08E-09 |
| BYSL | ENSG00000112578 | 0,390713 | 3,44E-09 |
| H4C14 | ENSG00000270882 | 1,024172 | 7,35E-09 |
| NECTIN4 | ENSG00000143217 | 1,041537 | 1,76E-08 |
| NFKBIL1 | ENSG00000204498 | 0,512233 | 3,66E-08 |
| PRKAB1 | ENSG00000111725 | 0,389225 | 7,55E-08 |
| EPB41L4A-AS1 | ENSG00000224032 | 0,442367 | 7,91E-08 |
| ENSG00000273199 | ENSG00000273199 | 1,169854 | 8,09E-08 |

| | | | |
|---|---|---|---|
| SLC3A2 | ENSG00000168003 | 0,403121 | 1,39E-07 |
| H2AW | ENSG00000181218 | 0,646834 | 1,57E-07 |
| GADD45A | ENSG00000116717 | 0,399395 | 1,94E-07 |
| PPFIA4 | ENSG00000143847 | -0,72668 | 4,26E-07 |
| ETV1 | ENSG00000006468 | 1,070357 | 5,40E-07 |
| RAD54L | ENSG00000085999 | 0,376355 | 7,13E-07 |
| BBC3 | ENSG00000105327 | 0,479891 | 1,20E-06 |
| TRMT61A | ENSG00000166166 | 0,376449 | 1,71E-06 |
| TRIAP1 | ENSG00000170855 | 0,373423 | 1,74E-06 |
| FDXR | ENSG00000161513 | 0,439896 | 2,02E-06 |
| GOLGA8B | ENSG00000215252 | -0,36941 | 2,02E-06 |
| ZNF764 | ENSG00000169951 | 0,365352 | 2,98E-06 |
| PPP1R15A | ENSG00000087074 | 0,354025 | 3,40E-06 |
| DUSP8 | ENSG00000184545 | 0,360598 | 4,73E-06 |
| PRR14 | ENSG00000156858 | 0,343673 | 4,93E-06 |
| CHTOP | ENSG00000160679 | 0,308596 | 7,20E-06 |
| ORMDL2 | ENSG00000123353 | 0,516127 | 9,96E-06 |
| USP30 | ENSG00000135093 | 0,419519 | 1,11E-05 |
| R3HDM2 | ENSG00000179912 | -0,40962 | 1,55E-05 |
| ENSG00000288999 | ENSG00000288999 | 0,700538 | 1,74E-05 |
| ZBTB17 | ENSG00000116809 | 0,417907 | 1,78E-05 |
| TOE1 | ENSG00000132773 | 0,346591 | 2,00E-05 |
| IP6K2 | ENSG00000068745 | 0,308371 | 2,02E-05 |
| H2BC12 | ENSG00000197903 | 0,493824 | 2,05E-05 |
| ZNF750 | ENSG00000141579 | 1,179761 | 2,05E-05 |
| ENSG00000285043 | ENSG00000285043 | 0,382207 | 2,11E-05 |
| LIF-AS2 | ENSG00000268812 | 1,290888 | 2,45E-05 |
| ENSG00000284602 | ENSG00000284602 | 0,769043 | 2,45E-05 |
| SORCS2 | ENSG00000184985 | 0,938784 | 2,48E-05 |
| PPP2R2A | ENSG00000221914 | 0,289468 | 2,85E-05 |
| DNAH3 | ENSG00000158486 | 1,485673 | 3,19E-05 |
| BRF2 | ENSG00000104221 | 0,429926 | 3,27E-05 |
| ZNF226 | ENSG00000167380 | 0,43574 | 3,27E-05 |
| FOS | ENSG00000170345 | -0,60171 | 3,65E-05 |
| RRP7BP | ENSG00000182841 | 0,350479 | 3,72E-05 |
| DINOL | ENSG00000285244 | 1,094767 | 3,94E-05 |
| ENSG00000285901 | ENSG00000285901 | 1,154259 | 3,94E-05 |
| PAN3-AS1 | ENSG00000261485 | 0,876738 | 4,03E-05 |
| NBEAL2 | ENSG00000160796 | -0,37157 | 4,59E-05 |
| CARS1 | ENSG00000110619 | 0,288033 | 4,76E-05 |
| ZNF593 | ENSG00000142684 | 0,416715 | 5,86E-05 |
| DUSP14 | ENSG00000276023 | 0,328549 | 5,89E-05 |
| NHLH2 | ENSG00000177551 | 0,533624 | 6,60E-05 |
| CDT1 | ENSG00000167513 | 0,348884 | 7,29E-05 |
| ZNF576 | ENSG00000124444 | 0,453363 | 7,52E-05 |
| CLK3 | ENSG00000179335 | 0,312259 | 8,72E-05 |
| JMJD7-PLA2G4B | ENSG00000168970 | -0,54861 | 8,74E-05 |

| | | | |
|---|---|---|---|
| FAM174A | ENSG00000174132 | 0,470722 | 0,000106 |
| ENSG00000276131 | ENSG00000276131 | 0,690424 | 0,000108 |
| H4C11 | ENSG00000197238 | 1,212138 | 0,00011 |
| PSMC3IP | ENSG00000131470 | 0,285472 | 0,00011 |
| SGPP2 | ENSG00000163082 | -0,57992 | 0,000113 |
| EPHA2 | ENSG00000142627 | 0,326389 | 0,000117 |
| NRAV | ENSG00000248008 | 0,38373 | 0,000123 |
| ENSG00000273391 | ENSG00000273391 | 0,796033 | 0,00014 |
| ZNF408 | ENSG00000175213 | 0,346128 | 0,00015 |
| LUC7L | ENSG00000007392 | 0,31215 | 0,000152 |
| ENSG00000260805 | ENSG00000260805 | 0,74051 | 0,000152 |
| RIBC2 | ENSG00000128408 | 0,55482 | 0,00017 |
| CCNB1IP1 | ENSG00000100814 | 0,313485 | 0,000178 |
| SLC38A2 | ENSG00000134294 | -0,35298 | 0,000178 |
| ENSG00000268403 | ENSG00000268403 | 0,53784 | 0,000178 |
| ZNF335 | ENSG00000198026 | 0,296094 | 0,000182 |
| ENSG00000230021 | ENSG00000230021 | -1,5406 | 0,000182 |
| DDX19B | ENSG00000157349 | 0,320524 | 0,000184 |
| AKIRIN2 | ENSG00000135334 | 0,284271 | 0,000185 |
| POMZP3 | ENSG00000146707 | 0,413644 | 0,000185 |
| RASSF1 | ENSG00000068028 | 0,302186 | 0,000186 |
| CLK1 | ENSG00000013441 | 0,282911 | 0,000204 |
| SNHG32 | ENSG00000204387 | 0,31402 | 0,000208 |
| ENSG00000271918 | ENSG00000271918 | 0,930686 | 0,000211 |
| MOK | ENSG00000080823 | 0,356168 | 0,000226 |
| C17orf97 | ENSG00000187624 | 0,424903 | 0,000235 |
| JPT1 | ENSG00000189159 | 0,293565 | 0,000239 |
| TIPIN | ENSG00000075131 | 0,302327 | 0,00024 |
| MUL1 | ENSG00000090432 | 0,306582 | 0,00024 |
| MED26 | ENSG00000105085 | 0,333149 | 0,000243 |
| DCAF16 | ENSG00000163257 | -0,31875 | 0,00027 |
| COL4A4 | ENSG00000081052 | -1,53039 | 0,00027 |
| PRCC | ENSG00000143294 | 0,291034 | 0,000271 |
| UBALD2 | ENSG00000185262 | 0,410239 | 0,000271 |
| TBC1D22A-DT | ENSG00000260708 | 0,538038 | 0,000312 |
| EMC9 | ENSG00000100908 | 0,362728 | 0,000321 |
| PDRG1 | ENSG00000088356 | 0,32951 | 0,000332 |
| NEB | ENSG00000183091 | -0,62418 | 0,000341 |
| BTNL9 | ENSG00000165810 | -0,45113 | 0,000342 |
| SLC16A9 | ENSG00000165449 | -0,38305 | 0,000366 |
| SLC12A4 | ENSG00000124067 | 0,440601 | 0,000389 |
| PRTG | ENSG00000166450 | -0,66188 | 0,000389 |
| ILF3-DT | ENSG00000267100 | 0,358948 | 0,000389 |
| AKAP8L | ENSG00000011243 | 0,302225 | 0,000403 |
| NTMT1 | ENSG00000148335 | 0,338057 | 0,000403 |
| AURKAP1 | ENSG00000213033 | 1,472529 | 0,000413 |
| ENSG00000286001 | ENSG00000286001 | 0,700855 | 0,000413 |

| | | | |
|---|---|---|---|
| CNOT3 | ENSG00000088038 | 0,299398 | 0,000425 |
| LENG1 | ENSG00000105617 | 0,502342 | 0,000442 |
| BUD31 | ENSG00000106245 | 0,326277 | 0,000449 |
| PDE4D | ENSG00000113448 | -0,5459 | 0,000449 |
| ENSG00000264112 | ENSG00000264112 | -0,36732 | 0,000449 |
| RBM4 | ENSG00000173933 | 0,273635 | 0,000457 |
| ENSG00000286116 | ENSG00000286116 | 1,078929 | 0,000457 |
| BANP | ENSG00000172530 | 0,33255 | 0,000463 |
| VPS13C | ENSG00000129003 | -0,41014 | 0,000463 |
| WDR46 | ENSG00000227057 | 0,275405 | 0,000531 |
| DUS3L | ENSG00000141994 | 0,307239 | 0,000551 |
| XRCC6P1 | ENSG00000237417 | 1,045026 | 0,000551 |
| CCDC174 | ENSG00000154781 | 0,338508 | 0,000557 |
| SOX4 | ENSG00000124766 | -0,26468 | 0,000563 |
| CHIC2 | ENSG00000109220 | 0,374727 | 0,000568 |
| LTB4R2 | ENSG00000213906 | -0,52568 | 0,000568 |
| LINC02846 | ENSG00000260193 | 0,72831 | 0,000591 |
| HROB | ENSG00000125319 | 0,345609 | 0,000594 |
| C1orf50 | ENSG00000164008 | 0,445824 | 0,000613 |
| ENSG00000283515 | ENSG00000283515 | 0,533379 | 0,000638 |
| CDIP1 | ENSG00000089486 | 0,330282 | 0,000646 |
| ENSG00000290058 | ENSG00000290058 | 1,195766 | 0,000724 |
| SNHG20 | ENSG00000234912 | 0,318458 | 0,000731 |
| GABARAPL1 | ENSG00000139112 | 0,304206 | 0,000754 |
| NLE1 | ENSG00000073536 | 0,26388 | 0,000755 |
| H6PD | ENSG00000049239 | -0,36578 | 0,000776 |
| ACADSB | ENSG00000196177 | -0,41993 | 0,000808 |
| GAS5 | ENSG00000234741 | 0,308511 | 0,000823 |
| INHBE | ENSG00000139269 | 0,751107 | 0,000864 |
| ZNHIT2 | ENSG00000174276 | 0,391963 | 0,000864 |
| RNF25 | ENSG00000163481 | 0,308139 | 0,000937 |
| THBS4 | ENSG00000113296 | -0,69314 | 0,000938 |
| GRIN2C | ENSG00000161509 | -0,47088 | 0,000994 |
| IP6K1 | ENSG00000176095 | 0,241743 | 0,000994 |
| ZSCAN21 | ENSG00000166529 | 0,335864 | 0,00101 |
| BPNT2 | ENSG00000104331 | -0,28762 | 0,001078 |
| EIF2B2 | ENSG00000119718 | 0,257215 | 0,001088 |
| CEP89 | ENSG00000121289 | 0,263358 | 0,001105 |
| WNK4 | ENSG00000126562 | -0,49572 | 0,001123 |
| GRWD1 | ENSG00000105447 | 0,270023 | 0,001155 |
| SERTAD1 | ENSG00000197019 | 0,505826 | 0,001155 |
| ZKSCAN2-DT | ENSG00000274925 | 0,476858 | 0,001165 |
| TRMT1 | ENSG00000104907 | 0,266229 | 0,001171 |
| CCDC86 | ENSG00000110104 | 0,270535 | 0,001171 |
| JUN | ENSG00000177606 | 0,281259 | 0,001171 |
| ENSG00000283235 | ENSG00000283235 | 1,157329 | 0,001177 |
| DTL | ENSG00000143476 | 0,313492 | 0,001178 |

| | | | |
|---|---|---|---|
| PGF | ENSG00000119630 | 0,475252 | 0,001213 |
| ATG101 | ENSG00000123395 | 0,33371 | 0,001213 |
| ENSG00000228793 | ENSG00000228793 | 1,239987 | 0,001213 |
| RAE1 | ENSG00000101146 | 0,238811 | 0,001215 |
| ZNF16 | ENSG00000170631 | 0,334769 | 0,001259 |
| ATM | ENSG00000149311 | -0,41835 | 0,001398 |
| DHX37 | ENSG00000150990 | 0,23646 | 0,001462 |
| SHANK2 | ENSG00000162105 | -0,9295 | 0,001577 |
| RPP38 | ENSG00000152464 | 0,288122 | 0,001619 |
| HSPA12A | ENSG00000165868 | -0,31739 | 0,001622 |
| MIR34AHG | ENSG00000228526 | 0,518119 | 0,001657 |
| LPP | ENSG00000145012 | -0,45874 | 0,001669 |
| IRS4 | ENSG00000133124 | -0,34527 | 0,001695 |
| ZNF689 | ENSG00000156853 | 0,259689 | 0,001873 |
| CAPN10-DT | ENSG00000260942 | 0,477122 | 0,001881 |
| SNHG4 | ENSG00000281398 | -0,29847 | 0,001959 |
| UBOX5 | ENSG00000185019 | 0,348924 | 0,002119 |
| ENSG00000269399 | ENSG00000269399 | 0,552349 | 0,002163 |
| RPS27L | ENSG00000185088 | 0,397545 | 0,002257 |
| PIBF1 | ENSG00000083535 | 0,25392 | 0,002291 |
| AHI1 | ENSG00000135541 | -0,35505 | 0,002319 |
| NR1D2 | ENSG00000174738 | -0,38141 | 0,002319 |
| ENSG00000228528 | ENSG00000228528 | 1,03971 | 0,00241 |
| ZNF48 | ENSG00000180035 | 0,260135 | 0,002541 |
| FIGNL1 | ENSG00000132436 | -0,31084 | 0,002558 |
| TP53I3 | ENSG00000115129 | 0,38593 | 0,002633 |
| NOP16 | ENSG00000048162 | 0,277671 | 0,002688 |
| PPP1R37 | ENSG00000104866 | 0,316027 | 0,002688 |
| REXO4 | ENSG00000148300 | 0,248608 | 0,002762 |
| TSSC4 | ENSG00000184281 | 0,299457 | 0,002762 |
| SRSF6 | ENSG00000124193 | 0,235398 | 0,002791 |
| UTP14A | ENSG00000156697 | 0,233565 | 0,002838 |
| ANKRD36BP1 | ENSG00000214262 | -0,87987 | 0,002843 |
| STK26 | ENSG00000134602 | -0,31689 | 0,002869 |
| ERO1A | ENSG00000197930 | -0,27111 | 0,002912 |
| PHF23 | ENSG00000040633 | 0,282053 | 0,002925 |
| LAMA3 | ENSG00000053747 | -0,28489 | 0,002952 |
| CCDC9 | ENSG00000105321 | 0,292043 | 0,002952 |
| FKBPL | ENSG00000204315 | 0,391876 | 0,002952 |
| LINC01311 | ENSG00000260924 | 0,474468 | 0,002952 |
| PLK2 | ENSG00000145632 | 0,460385 | 0,003083 |
| UNC5B | ENSG00000107731 | -0,24304 | 0,003117 |
| AARSD1 | ENSG00000266967 | 0,337894 | 0,003125 |
| XRCC1 | ENSG00000073050 | 0,24732 | 0,003181 |
| GLS | ENSG00000115419 | -0,35587 | 0,003186 |
| BRIP1 | ENSG00000136492 | -0,33905 | 0,003186 |
| BAK1 | ENSG00000030110 | 0,26195 | 0,003295 |

| | | | |
|---|---|---|---|
| TTLL3 | ENSG00000214021 | -0,26417 | 0,003371 |
| KLF13 | ENSG00000169926 | -0,37235 | 0,003409 |
| LINC02086 | ENSG00000244649 | 0,653262 | 0,003409 |
| TRIT1 | ENSG00000043514 | 0,238656 | 0,003436 |
| SLC16A8 | ENSG00000100156 | -1,28676 | 0,003452 |
| POFUT2 | ENSG00000186866 | 0,239733 | 0,003456 |
| PARTICL | ENSG00000286532 | 0,508414 | 0,003458 |
| MFSD2A | ENSG00000168389 | 0,428111 | 0,003472 |
| SNX10 | ENSG00000086300 | -0,44387 | 0,003539 |
| PKD1 | ENSG00000008710 | -0,30266 | 0,003655 |
| ZFAS1 | ENSG00000177410 | 0,338366 | 0,003655 |
| AAR2 | ENSG00000131043 | 0,249691 | 0,003756 |
| KMT2E-AS1 | ENSG00000239569 | 0,764497 | 0,003756 |
| NOL12 | ENSG00000273899 | 0,294788 | 0,003828 |
| PCBP1 | ENSG00000169564 | 0,238713 | 0,004036 |
| MARS1 | ENSG00000166986 | 0,222869 | 0,004074 |
| ENSG00000276791 | ENSG00000276791 | 0,509536 | 0,00415 |
| WRAP53 | ENSG00000141499 | 0,291763 | 0,004177 |
| LINC02983 | ENSG00000234432 | 0,647912 | 0,004177 |
| EGR1 | ENSG00000120738 | -0,40748 | 0,004267 |
| LTBP2 | ENSG00000119681 | -0,48372 | 0,004311 |
| GALNT8 | ENSG00000130035 | 0,959409 | 0,004372 |
| SMIM14 | ENSG00000163683 | -0,7476 | 0,004394 |
| ZNF473 | ENSG00000142528 | 0,26376 | 0,00448 |
| LDLRAD4 | ENSG00000168675 | -0,79879 | 0,00448 |
| RRN3P1 | ENSG00000248124 | 0,301258 | 0,00448 |
| ENSG00000257176 | ENSG00000257176 | -0,76582 | 0,00448 |
| TARBP1 | ENSG00000059588 | -0,26993 | 0,004639 |
| DOLPP1 | ENSG00000167130 | 0,271565 | 0,004639 |
| PVR | ENSG00000073008 | 0,236047 | 0,004711 |
| MRM2 | ENSG00000122687 | 0,24005 | 0,004866 |
| ENSG00000283341 | ENSG00000283341 | 0,600506 | 0,004866 |
| PDZK1 | ENSG00000174827 | 0,920861 | 0,004887 |
| ENSG00000280211 | ENSG00000280211 | 0,264836 | 0,004889 |
| CRNKL1 | ENSG00000101343 | 0,283614 | 0,00537 |
| PSMC3 | ENSG00000165916 | 0,246153 | 0,00537 |
| KTI12 | ENSG00000198841 | 0,288973 | 0,005434 |
| VPS37B | ENSG00000139722 | 0,229805 | 0,005575 |
| GTSE1-DT | ENSG00000277232 | 1,175864 | 0,005654 |
| ACAP2 | ENSG00000114331 | -0,43125 | 0,005698 |
| CCDC59 | ENSG00000133773 | 0,249849 | 0,00571 |
| C14orf119 | ENSG00000179933 | 0,266061 | 0,005763 |
| LINC00641 | ENSG00000258441 | -0,2501 | 0,005767 |
| ADCY7 | ENSG00000121281 | -0,67495 | 0,005804 |
| POP7 | ENSG00000172336 | 0,281556 | 0,005897 |
| SCAMP1-AS1 | ENSG00000245556 | 0,411795 | 0,005897 |
| ENSG00000269937 | ENSG00000269937 | -0,47762 | 0,005897 |

| | | | |
|---|---|---|---|
| SLC25A25 | ENSG00000148339 | 0,240452 | 0,005952 |
| PAF1 | ENSG00000006712 | 0,216743 | 0,006041 |
| PMS2P3 | ENSG00000127957 | 0,338344 | 0,006041 |
| RBMS1 | ENSG00000153250 | -0,31467 | 0,006041 |
| ZNF513 | ENSG00000163795 | 0,35204 | 0,006041 |
| ZNF581 | ENSG00000171425 | 0,289073 | 0,006041 |
| KCTD12 | ENSG00000178695 | -0,30136 | 0,006041 |
| SCFD2 | ENSG00000184178 | 0,294263 | 0,006041 |
| ZDHHC23 | ENSG00000184307 | -0,3897 | 0,006041 |
| SH3BP5-AS1 | ENSG00000224660 | -0,42356 | 0,006041 |
| PAPOLA-DT | ENSG00000260806 | 0,615697 | 0,006041 |
| ENSG00000289523 | ENSG00000289523 | 0,928028 | 0,006045 |
| HSD17B7P2 | ENSG00000099251 | 0,507013 | 0,006064 |
| MRPL22 | ENSG00000082515 | 0,277192 | 0,006087 |
| BIVM | ENSG00000134897 | -0,34568 | 0,006104 |
| CLASRP | ENSG00000104859 | 0,236153 | 0,006181 |
| TIMM9 | ENSG00000100575 | 0,25922 | 0,006184 |
| ZNF503-AS2 | ENSG00000237149 | 0,322775 | 0,006229 |
| RCN2 | ENSG00000117906 | -0,24656 | 0,006247 |
| DNHD1 | ENSG00000179532 | -0,46634 | 0,006346 |
| ENSG00000288983 | ENSG00000288983 | 0,368361 | 0,006381 |
| DDX39A | ENSG00000123136 | 0,217014 | 0,006391 |
| SCAND1 | ENSG00000171222 | 0,373003 | 0,006411 |
| KANK3 | ENSG00000186994 | 0,555349 | 0,006411 |
| RETREG1 | ENSG00000154153 | -0,47007 | 0,006465 |
| TK1 | ENSG00000167900 | 0,267839 | 0,006503 |
| KNTC1 | ENSG00000184445 | -0,23286 | 0,006647 |
| MRTO4 | ENSG00000053372 | 0,269145 | 0,006699 |
| BRIX1 | ENSG00000113460 | 0,220797 | 0,006699 |
| WDR31 | ENSG00000148225 | -0,4231 | 0,006713 |
| ZNF205 | ENSG00000122386 | 0,281593 | 0,006852 |
| TBRG4 | ENSG00000136270 | 0,234303 | 0,006852 |
| C18orf21 | ENSG00000141428 | 0,252107 | 0,006852 |
| C19orf54 | ENSG00000188493 | 0,25139 | 0,006852 |
| CTH | ENSG00000116761 | 0,249267 | 0,006859 |
| HMCN1 | ENSG00000143341 | -0,93631 | 0,006972 |
| COL5A2 | ENSG00000204262 | -0,43452 | 0,006972 |
| SARNP | ENSG00000205323 | 0,275617 | 0,006972 |
| CDADC1 | ENSG00000102543 | 0,411554 | 0,007037 |
| ZNF584 | ENSG00000171574 | 0,299525 | 0,007105 |
| KATNAL1 | ENSG00000102781 | -0,4174 | 0,007147 |
| ACSBG1 | ENSG00000103740 | 0,323067 | 0,007147 |
| NXT1 | ENSG00000132661 | 0,327959 | 0,007147 |
| OSER1 | ENSG00000132823 | 0,290424 | 0,007147 |
| DNAH17-AS1 | ENSG00000267432 | 0,925733 | 0,007147 |
| CARNMT1 | ENSG00000156017 | -0,31493 | 0,007199 |
| PPP2R3C | ENSG00000092020 | 0,260481 | 0,007204 |

| | | | |
|---|---|---|---|
| THAP7 | ENSG00000184436 | 0,240414 | 0,00722 |
| EIF1 | ENSG00000173812 | 0,244677 | 0,007261 |
| SLC44A5 | ENSG00000137968 | -0,37014 | 0,007298 |
| LINC01004 | ENSG00000228393 | 0,467756 | 0,007309 |
| SNRPA | ENSG00000077312 | 0,264443 | 0,007396 |
| KLHL26 | ENSG00000167487 | 0,302934 | 0,007535 |
| SFPQ | ENSG00000116560 | 0,242328 | 0,007539 |
| ZBTB7B | ENSG00000160685 | 0,276851 | 0,007579 |
| ZNF672 | ENSG00000171161 | 0,256922 | 0,007579 |
| CCDC137 | ENSG00000185298 | 0,232705 | 0,007579 |
| TIGD1 | ENSG00000221944 | 0,287035 | 0,007662 |
| BRPF1 | ENSG00000156983 | 0,222282 | 0,007685 |
| ENSG00000289043 | ENSG00000289043 | 0,542721 | 0,007685 |
| TMSB15B | ENSG00000158427 | 1,362735 | 0,007715 |
| SARS1 | ENSG00000031698 | 0,211112 | 0,00787 |
| H2AC19 | ENSG00000288859 | 2,703408 | 0,00787 |
| GALNT7 | ENSG00000109586 | -0,36162 | 0,007969 |
| EIF1B | ENSG00000114784 | 0,236189 | 0,007969 |
| IQCC | ENSG00000160051 | 0,240718 | 0,007984 |
| SLC25A19 | ENSG00000125454 | 0,265623 | 0,007996 |
| NIFK | ENSG00000155438 | 0,236206 | 0,007996 |
| COA6 | ENSG00000168275 | 0,260567 | 0,007996 |
| ZNF320 | ENSG00000182986 | -0,53621 | 0,008027 |
| H2BU1 | ENSG00000196890 | 0,858979 | 0,008229 |
| SUCO | ENSG00000094975 | -0,32931 | 0,008289 |
| RABGEF1P1 | ENSG00000229180 | 0,278414 | 0,008358 |
| ALDH5A1 | ENSG00000112294 | -0,28761 | 0,008709 |
| GOLGA6B | ENSG00000215186 | -1,28013 | 0,008709 |
| TOM1 | ENSG00000100284 | 0,280659 | 0,008723 |
| ERBIN | ENSG00000112851 | -0,29482 | 0,008723 |
| HIRIP3 | ENSG00000149929 | 0,233266 | 0,008723 |
| C5orf24 | ENSG00000181904 | -0,25661 | 0,008723 |
| CNPY2-AS1 | ENSG00000257303 | 0,503043 | 0,008723 |
| CDC42EP1 | ENSG00000128283 | 0,265209 | 0,008772 |
| ZNF337 | ENSG00000130684 | 0,246998 | 0,008772 |
| RUFY1 | ENSG00000176783 | 0,209687 | 0,008772 |
| MZT1 | ENSG00000204899 | -0,2477 | 0,008772 |
| TCF4 | ENSG00000196628 | -0,40368 | 0,008861 |
| NME1-NME2 | ENSG00000011052 | 0,377792 | 0,008912 |
| H2BC21 | ENSG00000184678 | 0,461292 | 0,008912 |
| RBPJ | ENSG00000168214 | -0,24009 | 0,009089 |
| TMEM11 | ENSG00000178307 | 0,283077 | 0,009089 |
| ELMOD2 | ENSG00000179387 | -0,27242 | 0,009089 |
| ARPIN-AP3S2 | ENSG00000250021 | 0,256742 | 0,009089 |
| CFAP43 | ENSG00000197748 | -0,5189 | 0,009288 |
| ENSG00000204666 | ENSG00000204666 | 0,90477 | 0,009313 |
| VWCE | ENSG00000167992 | -0,48951 | 0,009342 |

| | | | |
|---|---|---|---|
| DNAH17 | ENSG00000187775 | 0,450284 | 0,009567 |
| KIAA1109 | ENSG00000138688 | -0,37014 | 0,009572 |
| CD109 | ENSG00000156535 | -0,51379 | 0,009572 |
| RAP1GAP2 | ENSG00000132359 | -0,31969 | 0,009577 |
| PSTK | ENSG00000179988 | 0,352835 | 0,009667 |
| BRCA2 | ENSG00000139618 | -0,28735 | 0,00986 |
| TPH1 | ENSG00000129167 | -0,56247 | 0,009908 |
| PPP2R2D | ENSG00000175470 | 0,243736 | 0,009995 |
| GEMIN8 | ENSG00000046647 | 0,313782 | 0,01028 |
| DNM1 | ENSG00000106976 | -0,28402 | 0,01028 |
| RSKR | ENSG00000167524 | -0,45679 | 0,01028 |
| FAM156B | ENSG00000179304 | -0,37894 | 0,010312 |
| ANK2 | ENSG00000145362 | -0,30404 | 0,010336 |
| CDK6 | ENSG00000105810 | -0,31361 | 0,010378 |
| NXPE3 | ENSG00000144815 | -0,35382 | 0,010378 |
| PNO1 | ENSG00000115946 | 0,207225 | 0,01038 |
| NFYC | ENSG00000066136 | 0,22609 | 0,010427 |
| LINC01341 | ENSG00000227953 | -0,75835 | 0,010427 |
| LRRC8B | ENSG00000197147 | -0,33583 | 0,010456 |
| EMB | ENSG00000170571 | -0,31373 | 0,010593 |
| SGTA | ENSG00000104969 | 0,226566 | 0,010625 |
| PLOD2 | ENSG00000152952 | -0,26531 | 0,010625 |
| INTS5 | ENSG00000185085 | 0,237023 | 0,010625 |
| TMEM123 | ENSG00000152558 | -0,27781 | 0,010655 |
| MRPL49 | ENSG00000149792 | 0,223349 | 0,010917 |
| CAP2 | ENSG00000112186 | -0,38939 | 0,011011 |
| SNHG30 | ENSG00000267321 | 0,360253 | 0,011168 |
| KIF14 | ENSG00000118193 | -0,31106 | 0,011234 |
| TRPM7 | ENSG00000092439 | -0,30117 | 0,011234 |
| MBNL2 | ENSG00000139793 | -0,51181 | 0,011314 |
| RNF125 | ENSG00000101695 | -0,34177 | 0,011378 |
| C19orf48 | ENSG00000167747 | 0,238463 | 0,011608 |
| ATP8A1 | ENSG00000124406 | -0,49521 | 0,011611 |
| TMED7 | ENSG00000134970 | -0,32721 | 0,011641 |
| DYNLT2 | ENSG00000184786 | 0,388377 | 0,011641 |
| ZNF117 | ENSG00000152926 | -0,53772 | 0,011769 |
| FAM157C | ENSG00000260528 | -0,59433 | 0,011825 |
| EIF3D | ENSG00000100353 | 0,221323 | 0,012068 |
| TAS2R5 | ENSG00000127366 | -0,98121 | 0,012068 |
| EDA2R | ENSG00000131080 | 0,580922 | 0,012068 |
| ALAS1 | ENSG00000023330 | 0,201064 | 0,012281 |
| SUPT7L | ENSG00000119760 | 0,21624 | 0,012281 |
| SNHG10 | ENSG00000247092 | 0,315995 | 0,012281 |
| AVPR1A | ENSG00000166148 | -0,81343 | 0,012314 |
| GEMIN7 | ENSG00000142252 | 0,270977 | 0,012388 |
| BLOC1S2 | ENSG00000196072 | 0,240015 | 0,012399 |
| ENSG00000289028 | ENSG00000289028 | 0,579406 | 0,012399 |

| | | | |
|---|---|---|---|
| TFPT | ENSG00000105619 | 0,343645 | 0,012514 |
| BCL7B | ENSG00000106635 | 0,236496 | 0,012514 |
| TMEM170B | ENSG00000205269 | -0,4092 | 0,012514 |
| UIMC1 | ENSG00000087206 | 0,212248 | 0,012543 |
| FMN2 | ENSG00000155816 | -0,25667 | 0,012653 |
| GOT1 | ENSG00000120053 | 0,245159 | 0,012745 |
| SLAIN2 | ENSG00000109171 | -0,40803 | 0,012783 |
| NIPBL-DT | ENSG00000285967 | 0,274428 | 0,012939 |
| RIOK1 | ENSG00000124784 | 0,197825 | 0,013009 |
| CGB7 | ENSG00000196337 | 0,774539 | 0,013216 |
| DPF2 | ENSG00000133884 | 0,200583 | 0,013292 |
| C2CD3 | ENSG00000168014 | 0,22869 | 0,013627 |
| SOX6 | ENSG00000110693 | -0,6633 | 0,013653 |
| SAR1B | ENSG00000152700 | 0,243601 | 0,013718 |
| ZFAT | ENSG00000066827 | 0,365054 | 0,013775 |
| RWDD1 | ENSG00000111832 | 0,255571 | 0,013775 |
| ZNF296 | ENSG00000170684 | 0,320651 | 0,013775 |
| PEA15 | ENSG00000162734 | 0,204473 | 0,013793 |
| IPO5P1 | ENSG00000269837 | 0,802385 | 0,013793 |
| TMEM115 | ENSG00000126062 | 0,232926 | 0,013842 |
| MAVS | ENSG00000088888 | -0,22334 | 0,013938 |
| NUP58 | ENSG00000139496 | -0,23306 | 0,01397 |
| KLF10 | ENSG00000155090 | -0,26041 | 0,01397 |
| FAM185A | ENSG00000222011 | 0,414009 | 0,01397 |
| OVGP1 | ENSG00000085465 | 0,251434 | 0,014143 |
| RPP38-DT | ENSG00000176236 | 0,615862 | 0,01423 |
| CDCA5 | ENSG00000146670 | 0,199035 | 0,014331 |
| VLDLR | ENSG00000147852 | -0,4826 | 0,014331 |
| ZNF574 | ENSG00000105732 | 0,245679 | 0,01444 |
| GOSR1 | ENSG00000108587 | 0,261434 | 0,014455 |
| BAX | ENSG00000087088 | 0,297215 | 0,014463 |
| SRSF3 | ENSG00000112081 | 0,206245 | 0,014511 |
| TBL1XR1 | ENSG00000177565 | -0,27717 | 0,014511 |
| BEND4 | ENSG00000188848 | -0,37792 | 0,014511 |
| DENND1B | ENSG00000213047 | -0,41083 | 0,014511 |
| GTF2F1 | ENSG00000125651 | 0,207439 | 0,014536 |
| ZNF622 | ENSG00000173545 | 0,263 | 0,014536 |
| ALKBH1 | ENSG00000100601 | 0,238132 | 0,014631 |
| PLEKHG4B | ENSG00000153404 | -0,6649 | 0,014845 |
| KCTD5 | ENSG00000167977 | 0,198624 | 0,014845 |
| ZNF248 | ENSG00000198105 | -0,31671 | 0,015008 |
| PHF5A | ENSG00000100410 | 0,231629 | 0,015017 |
| OLIG2 | ENSG00000205927 | 0,287031 | 0,015017 |
| N6AMT1 | ENSG00000156239 | 0,246672 | 0,015043 |
| HLF | ENSG00000108924 | -0,47537 | 0,015046 |
| YY1AP1 | ENSG00000163374 | 0,194502 | 0,015199 |
| GMEB1 | ENSG00000162419 | 0,232091 | 0,015236 |

| | | | |
|---|---|---|---|
| RIMS3 | ENSG00000117016 | -0,37648 | 0,015338 |
| NCKAP1 | ENSG00000061676 | -0,21083 | 0,015419 |
| MRPS31 | ENSG00000102738 | 0,229673 | 0,015419 |
| ITGA2 | ENSG00000164171 | -0,48906 | 0,015419 |
| FGD6 | ENSG00000180263 | -0,48966 | 0,015419 |
| ZNF787 | ENSG00000142409 | 0,267247 | 0,015441 |
| ABCB10 | ENSG00000135776 | -0,24869 | 0,015517 |
| PSMG3 | ENSG00000157778 | 0,270185 | 0,015517 |
| PBDC1 | ENSG00000102390 | 0,32257 | 0,015599 |
| ZNF628 | ENSG00000197483 | 0,444885 | 0,015601 |
| CDK9 | ENSG00000136807 | 0,261949 | 0,015633 |
| SAC3D1 | ENSG00000168061 | 0,305843 | 0,015633 |
| AKAP5 | ENSG00000179841 | -0,46137 | 0,015633 |
| GOLGA8A | ENSG00000175265 | -0,25204 | 0,015661 |
| MDM1 | ENSG00000111554 | -0,27853 | 0,015723 |
| THAP3 | ENSG00000041988 | 0,281487 | 0,015727 |
| CFAP97 | ENSG00000164323 | -0,35184 | 0,015806 |
| OIP5-AS1 | ENSG00000247556 | -0,23337 | 0,015898 |
| ECHDC2 | ENSG00000121310 | -0,38328 | 0,015947 |
| HRAS | ENSG00000174775 | 0,276235 | 0,015963 |
| SEMA3A | ENSG00000075213 | -0,57237 | 0,016167 |
| GPATCH4 | ENSG00000160818 | 0,220279 | 0,016167 |
| AGAP5 | ENSG00000172650 | -0,32047 | 0,016167 |
| LMAN2L | ENSG00000114988 | 0,232699 | 0,016211 |
| GMNN | ENSG00000112312 | 0,208483 | 0,016249 |
| TRPS1 | ENSG00000104447 | -0,40587 | 0,016258 |
| DRG1 | ENSG00000185721 | 0,219661 | 0,016258 |
| C19orf47 | ENSG00000160392 | 0,294203 | 0,016425 |
| OAZ2 | ENSG00000180304 | 0,215402 | 0,016425 |
| OXLD1 | ENSG00000204237 | 0,290898 | 0,01657 |
| AUP1 | ENSG00000115307 | 0,212277 | 0,016617 |
| PCYOX1 | ENSG00000116005 | -0,26411 | 0,016617 |
| GRHL3 | ENSG00000158055 | 0,554822 | 0,016702 |
| CCT6P1 | ENSG00000228409 | 0,220377 | 0,016926 |
| LRP1 | ENSG00000123384 | -0,26704 | 0,017014 |
| ZNF385A | ENSG00000161642 | 0,338959 | 0,017122 |
| ANGEL2 | ENSG00000174606 | -0,23881 | 0,017163 |
| PRR7 | ENSG00000131188 | 0,314896 | 0,017262 |
| ZNF773 | ENSG00000152439 | 0,301003 | 0,017262 |
| MAGEF1 | ENSG00000177383 | 0,205342 | 0,017266 |
| WASHC4 | ENSG00000136051 | -0,31488 | 0,017301 |
| ZNF282 | ENSG00000170265 | 0,203478 | 0,017319 |
| ZRANB2-DT | ENSG00000229956 | -1,05546 | 0,017482 |
| ZNF768 | ENSG00000169957 | 0,224509 | 0,017526 |
| ZDHHC21 | ENSG00000175893 | -0,42946 | 0,017526 |
| METTL2A | ENSG00000087995 | 0,217191 | 0,017691 |
| RPS19BP1 | ENSG00000187051 | 0,251508 | 0,017775 |

| | | | |
|---|---|---|---|
| YARS1 | ENSG00000134684 | 0,200849 | 0,017896 |
| LRRC8C | ENSG00000171488 | -0,32194 | 0,017896 |
| SNAP47 | ENSG00000143740 | 0,218388 | 0,018053 |
| GBF1 | ENSG00000107862 | 0,190138 | 0,018119 |
| MANEA | ENSG00000172469 | -0,39899 | 0,018119 |
| TRIB1 | ENSG00000173334 | -0,28116 | 0,018119 |
| GVQW3 | ENSG00000179240 | -0,32967 | 0,018119 |
| LRRC66 | ENSG00000188993 | -0,71233 | 0,018119 |
| MASP2 | ENSG00000009724 | -0,56729 | 0,018124 |
| ARSB | ENSG00000113273 | -0,3361 | 0,018124 |
| DMD | ENSG00000198947 | -0,35775 | 0,018257 |
| CYHR1 | ENSG00000187954 | 0,218027 | 0,018444 |
| TMEM47 | ENSG00000147027 | -0,36747 | 0,018575 |
| FREM2 | ENSG00000150893 | -0,49473 | 0,018575 |
| ADAM9 | ENSG00000168615 | -0,2944 | 0,018575 |
| TMEM79 | ENSG00000163472 | 0,289167 | 0,018642 |
| MED19 | ENSG00000156603 | 0,317699 | 0,018761 |
| ZFAND3 | ENSG00000156639 | 0,21291 | 0,018966 |
| ENSG00000285608 | ENSG00000285608 | -0,32279 | 0,019055 |
| FUZ | ENSG00000010361 | 0,240079 | 0,019126 |
| ACER3 | ENSG00000078124 | -0,29413 | 0,019126 |
| SDHAF1 | ENSG00000205138 | 0,328404 | 0,019126 |
| FRG1 | ENSG00000109536 | 0,26173 | 0,019247 |
| CTNNBL1 | ENSG00000132792 | 0,279781 | 0,019249 |
| ADD3 | ENSG00000148700 | -0,30775 | 0,01942 |
| MRPL48 | ENSG00000175581 | 0,259823 | 0,01942 |
| ID4 | ENSG00000172201 | -0,20802 | 0,019433 |
| SPPL2B | ENSG00000005206 | -0,20667 | 0,019529 |
| INTS2 | ENSG00000108506 | -0,27511 | 0,019529 |
| FAM217B | ENSG00000196227 | -0,33403 | 0,019569 |
| NDUFC2-KCTD14 | ENSG00000259112 | 0,937007 | 0,019689 |
| SGF29 | ENSG00000176476 | 0,287366 | 0,019834 |
| BAG1 | ENSG00000107262 | 0,242454 | 0,019886 |
| SLC4A7 | ENSG00000033867 | -0,48388 | 0,019892 |
| PRR11 | ENSG00000068489 | -0,23298 | 0,019892 |
| MLLT10 | ENSG00000078403 | -0,244 | 0,019892 |
| MCOLN3 | ENSG00000055732 | -0,32895 | 0,019937 |
| TARS1 | ENSG00000113407 | 0,199996 | 0,019978 |
| ZNF711 | ENSG00000147180 | -0,27443 | 0,019978 |
| BCDIN3D | ENSG00000186666 | 0,284368 | 0,019978 |
| RRP8 | ENSG00000132275 | 0,220465 | 0,020033 |
| SPATA2 | ENSG00000158480 | 0,256672 | 0,020132 |
| CHPF2 | ENSG00000033100 | 0,202674 | 0,020438 |
| ANK3 | ENSG00000151150 | -0,26617 | 0,020439 |
| ZNF256 | ENSG00000152454 | 0,319781 | 0,020439 |
| NSMCE2 | ENSG00000156831 | 0,270236 | 0,020439 |
| FCHO2 | ENSG00000157107 | -0,39976 | 0,020439 |

| | | | |
|---|---|---|---|
| SRR | ENSG00000167720 | -0,40038 | 0,020439 |
| ICE2 | ENSG00000128915 | -0,22614 | 0,020899 |
| MAP1S | ENSG00000130479 | 0,215372 | 0,020899 |
| ZNF394 | ENSG00000160908 | 0,238226 | 0,020913 |
| ZDBF2 | ENSG00000204186 | -0,40076 | 0,020953 |
| SLC7A2 | ENSG00000003989 | -0,33046 | 0,020972 |
| DUSP12 | ENSG00000081721 | 0,249084 | 0,020972 |
| FAM20B | ENSG00000116199 | -0,232 | 0,020972 |
| RP9 | ENSG00000164610 | 0,267877 | 0,020972 |
| GPRASP1 | ENSG00000198932 | -0,6183 | 0,020972 |
| SCNM1 | ENSG00000163156 | 0,258478 | 0,021206 |
| DYNLT3 | ENSG00000165169 | -0,25871 | 0,021267 |
| TRMO | ENSG00000136932 | 0,289473 | 0,021275 |
| MKLN1 | ENSG00000128585 | 0,24717 | 0,021339 |
| SAMD5 | ENSG00000203727 | -0,48768 | 0,021339 |
| LINC01719 | ENSG00000233396 | -0,41213 | 0,021339 |
| TRAM1 | ENSG00000067167 | -0,2156 | 0,021419 |
| FBXL12 | ENSG00000127452 | 0,23865 | 0,021419 |
| HOXD10 | ENSG00000128710 | -0,23387 | 0,021419 |
| RRS1 | ENSG00000179041 | 0,203689 | 0,021419 |
| CSF1 | ENSG00000184371 | 0,295437 | 0,021505 |
| GP1BA | ENSG00000185245 | -0,61839 | 0,021505 |
| COA4 | ENSG00000181924 | 0,210927 | 0,021512 |
| PDE5A | ENSG00000138735 | -0,40394 | 0,021626 |
| CBWD5 | ENSG00000147996 | 0,229466 | 0,021626 |
| ENSG00000288061 | ENSG00000288061 | 0,816862 | 0,021996 |
| SMAD9 | ENSG00000120693 | -0,30697 | 0,022149 |
| ITPA | ENSG00000125877 | 0,250609 | 0,022149 |
| WDR4 | ENSG00000160193 | 0,200727 | 0,022149 |
| BIRC5 | ENSG00000089685 | 0,199682 | 0,022182 |
| SLC26A2 | ENSG00000155850 | -0,31418 | 0,022182 |
| LYPLA1 | ENSG00000120992 | -0,28238 | 0,022235 |
| ATXN7L2 | ENSG00000162650 | 0,290775 | 0,022235 |
| ZDHHC20 | ENSG00000180776 | -0,27782 | 0,022235 |
| MAD1L1 | ENSG00000002822 | 0,28566 | 0,022269 |
| PML | ENSG00000140464 | 0,245431 | 0,022269 |
| CAMK1D | ENSG00000183049 | -0,34858 | 0,022272 |
| STXBP3 | ENSG00000116266 | -0,25967 | 0,022416 |
| FAM161A | ENSG00000170264 | -0,38435 | 0,022416 |
| RNF213 | ENSG00000173821 | -0,24984 | 0,022416 |
| ADPRM | ENSG00000170222 | 0,351184 | 0,022588 |
| CHD1-DT | ENSG00000248489 | 0,59656 | 0,022588 |
| PM20D2 | ENSG00000146281 | -0,25441 | 0,022704 |
| DYM | ENSG00000141627 | 0,251257 | 0,022712 |
| DDX41 | ENSG00000183258 | 0,19183 | 0,022712 |
| PMS1 | ENSG00000064933 | -0,23091 | 0,02313 |
| LINC00342 | ENSG00000232931 | -0,33064 | 0,02313 |

| | | | |
|---|---|---|---|
| ENSG00000289021 | ENSG00000289021 | 0,498434 | 0,023185 |
| ARIH2OS | ENSG00000221883 | 0,499543 | 0,023191 |
| ESS2 | ENSG00000100056 | 0,232394 | 0,023193 |
| HOOK3 | ENSG00000168172 | -0,42809 | 0,023193 |
| LYSMD3 | ENSG00000176018 | -0,3341 | 0,023193 |
| SRPK3 | ENSG00000184343 | -0,80157 | 0,023304 |
| MID1 | ENSG00000101871 | -0,29345 | 0,023327 |
| ENSG00000286207 | ENSG00000286207 | -0,73908 | 0,023327 |
| RRP12 | ENSG00000052749 | 0,192383 | 0,023374 |
| TACO1 | ENSG00000136463 | 0,224636 | 0,023374 |
| DOK3 | ENSG00000146094 | -0,3992 | 0,023374 |
| RPS2P55 | ENSG00000216866 | 0,212283 | 0,023374 |
| SHMT2 | ENSG00000182199 | 0,195147 | 0,023425 |
| SORBS1 | ENSG00000095637 | 0,370878 | 0,023516 |
| RHBDD2 | ENSG00000005486 | 0,230316 | 0,023855 |
| DLC1 | ENSG00000164741 | -0,26925 | 0,023855 |
| RABGGTA | ENSG00000100949 | 0,28091 | 0,024032 |
| TBX15 | ENSG00000092607 | 0,618312 | 0,024047 |
| BORCS5 | ENSG00000165714 | 0,414675 | 0,024047 |
| NEDD4 | ENSG00000069869 | -0,34582 | 0,024164 |
| FEN1 | ENSG00000168496 | 0,195197 | 0,024235 |
| CSTF1 | ENSG00000101138 | 0,211171 | 0,024312 |
| MRPL27 | ENSG00000108826 | 0,235842 | 0,024312 |
| TMEM234 | ENSG00000160055 | 0,283736 | 0,024312 |
| SDHAP1 | ENSG00000185485 | 0,270886 | 0,024312 |
| ENSG00000273204 | ENSG00000273204 | 0,788672 | 0,024312 |
| ENSG00000260855 | ENSG00000260855 | -0,94155 | 0,024373 |
| SPRED2 | ENSG00000198369 | 0,355277 | 0,024519 |
| ENSG00000240399 | ENSG00000240399 | 0,889879 | 0,024681 |
| PFDN1 | ENSG00000113068 | 0,208013 | 0,02476 |
| C16orf91 | ENSG00000174109 | 0,299699 | 0,024963 |
| SLC39A7 | ENSG00000112473 | 0,194515 | 0,024988 |
| SEPTIN11 | ENSG00000138758 | -0,22942 | 0,024988 |
| SPA17 | ENSG00000064199 | 0,367728 | 0,025047 |
| SLX1A-SULT1A3 | ENSG00000213599 | -0,26856 | 0,025049 |
| ATP6V1D | ENSG00000100554 | 0,2249 | 0,025206 |
| SCOC | ENSG00000153130 | -0,23311 | 0,02524 |
| EMX2OS | ENSG00000229847 | 0,413571 | 0,02524 |
| RPP30 | ENSG00000148688 | 0,202834 | 0,025253 |
| CBX7 | ENSG00000100307 | -0,40148 | 0,025369 |
| PLCE1 | ENSG00000138193 | -0,28403 | 0,025442 |
| ZNRD2 | ENSG00000173465 | 0,256953 | 0,025482 |
| PLEKHA1 | ENSG00000107679 | -0,29405 | 0,025655 |
| HGSNAT | ENSG00000165102 | -0,24103 | 0,025655 |
| ZNF521 | ENSG00000198795 | -0,37176 | 0,025655 |
| STAM-DT | ENSG00000260589 | 0,381018 | 0,025729 |
| LMBR1 | ENSG00000105983 | -0,27349 | 0,025754 |

| | | | |
|---|---|---|---|
| SAAL1 | ENSG00000166788 | 0,204492 | 0,025754 |
| FAM126A | ENSG00000122591 | -0,28342 | 0,025785 |
| RPF2 | ENSG00000197498 | 0,202916 | 0,025785 |
| YBX3 | ENSG00000060138 | 0,198243 | 0,02583 |
| ZNF165 | ENSG00000197279 | 0,359153 | 0,025928 |
| SBF2 | ENSG00000133812 | 0,34525 | 0,025956 |
| FHIP2A | ENSG00000151553 | -0,28846 | 0,026176 |
| TMED8 | ENSG00000100580 | -0,25617 | 0,026217 |
| ZNF324 | ENSG00000083812 | 0,239413 | 0,026275 |
| USP39 | ENSG00000168883 | 0,183042 | 0,026363 |
| CPOX | ENSG00000080819 | -0,24022 | 0,026377 |
| LLPH | ENSG00000139233 | 0,232376 | 0,0266 |
| ENSG00000280383 | ENSG00000280383 | 0,540419 | 0,026611 |
| PTPRS | ENSG00000105426 | -0,27668 | 0,026707 |
| CACNA2D1 | ENSG00000153956 | -0,43954 | 0,026707 |
| ZMAT3 | ENSG00000172667 | 0,293641 | 0,026707 |
| CHDH | ENSG00000016391 | -0,25681 | 0,02714 |
| SRSF4 | ENSG00000116350 | 0,193915 | 0,027228 |
| PPM1D | ENSG00000170836 | 0,202784 | 0,027228 |
| ARL10 | ENSG00000175414 | -0,22653 | 0,027228 |
| ENSG00000256966 | ENSG00000256966 | -0,83117 | 0,027263 |
| UNG | ENSG00000076248 | 0,179863 | 0,027334 |
| SYT16 | ENSG00000139973 | -0,66071 | 0,027334 |
| MRPL36 | ENSG00000171421 | 0,28263 | 0,027334 |
| SLC39A10 | ENSG00000196950 | -0,24652 | 0,02767 |
| KRCC1 | ENSG00000172086 | -0,29055 | 0,027869 |
| PTER | ENSG00000165983 | -0,24116 | 0,028001 |
| PIH1D1 | ENSG00000104872 | 0,251823 | 0,028187 |
| INTS12 | ENSG00000138785 | 0,202133 | 0,028236 |
| RNF207 | ENSG00000158286 | -0,48621 | 0,028236 |
| FAS | ENSG00000026103 | 0,279896 | 0,028395 |
| DPY19L4 | ENSG00000156162 | -0,40907 | 0,028412 |
| WDR13 | ENSG00000101940 | 0,328715 | 0,028507 |
| ATP9A | ENSG00000054793 | -0,26689 | 0,02862 |
| AGFG1 | ENSG00000173744 | 0,218558 | 0,028622 |
| MRTFA | ENSG00000196588 | 0,223207 | 0,028638 |
| ITGA6 | ENSG00000091409 | -0,34429 | 0,028718 |
| C5 | ENSG00000106804 | -0,5499 | 0,028718 |
| MIB1 | ENSG00000101752 | -0,34071 | 0,028872 |
| TEX15 | ENSG00000133863 | -0,40893 | 0,028872 |
| DDX56 | ENSG00000136271 | 0,199202 | 0,028872 |
| MED8 | ENSG00000159479 | 0,211074 | 0,028872 |
| AP2A2 | ENSG00000183020 | 0,239851 | 0,028872 |
| POU2F1-DT | ENSG00000272205 | 0,578472 | 0,028888 |
| VPS25 | ENSG00000131475 | 0,202577 | 0,029089 |
| THG1L | ENSG00000113272 | 0,257226 | 0,029279 |
| PRRC1 | ENSG00000164244 | -0,25013 | 0,029566 |

| | | | |
|---|---|---|---|
| AIMP2 | ENSG00000106305 | 0,192203 | 0,029743 |
| NUP210 | ENSG00000132182 | -0,17882 | 0,029743 |
| EEFSEC | ENSG00000132394 | 0,24346 | 0,029743 |
| ACTR2 | ENSG00000138071 | -0,23264 | 0,029743 |
| BRAF | ENSG00000157764 | 0,285688 | 0,029743 |
| KREMEN1 | ENSG00000183762 | -0,22655 | 0,029743 |
| ENSG00000272906 | ENSG00000272906 | 0,672895 | 0,029743 |
| STX1B | ENSG00000099365 | -0,67687 | 0,029873 |
| ZFP30 | ENSG00000120784 | -0,2998 | 0,029873 |
| GPN2 | ENSG00000142751 | 0,218264 | 0,029873 |
| FBXO41 | ENSG00000163013 | -0,30325 | 0,029873 |
| ITGA1 | ENSG00000213949 | -0,49256 | 0,029873 |
| AKAP6 | ENSG00000151320 | -0,65287 | 0,029926 |
| BNIP1 | ENSG00000113734 | 0,239435 | 0,030177 |
| TMEM65 | ENSG00000164983 | -0,24859 | 0,030182 |
| LMAN1 | ENSG00000074695 | -0,22832 | 0,030242 |
| CCNB3 | ENSG00000147082 | -0,61245 | 0,030669 |
| SENP3-EIF4A1 | ENSG00000277957 | 0,337952 | 0,030838 |
| IL6ST | ENSG00000134352 | -0,31701 | 0,030937 |
| PAQR3 | ENSG00000163291 | -0,25824 | 0,030973 |
| SLFN5 | ENSG00000166750 | -0,60072 | 0,030973 |
| PUF60 | ENSG00000179950 | 0,195984 | 0,031005 |
| WASH5P | ENSG00000282458 | -0,32368 | 0,031192 |
| NEBL | ENSG00000078114 | -0,3947 | 0,031283 |
| PI4KB | ENSG00000143393 | 0,181461 | 0,031285 |
| MOAP1 | ENSG00000165943 | 0,207959 | 0,031285 |
| MIDEAS | ENSG00000156030 | -0,29677 | 0,031577 |
| UTP3 | ENSG00000132467 | 0,191932 | 0,031598 |
| MKX | ENSG00000150051 | -0,41742 | 0,031598 |
| WDR73 | ENSG00000177082 | 0,18515 | 0,031598 |
| MUC19 | ENSG00000205592 | 0,85523 | 0,031598 |
| ENSG00000289298 | ENSG00000289298 | 0,704287 | 0,031767 |
| PI4K2B | ENSG00000038210 | -0,2796 | 0,031982 |
| TCFL5 | ENSG00000101190 | -0,26329 | 0,032163 |
| PASK | ENSG00000115687 | -0,2004 | 0,032163 |
| DCTN5 | ENSG00000166847 | 0,205654 | 0,032163 |
| NUDT3 | ENSG00000272325 | -0,20983 | 0,032163 |
| SEL1L | ENSG00000071537 | -0,33665 | 0,032167 |
| FRK | ENSG00000111816 | -0,66677 | 0,032167 |
| NOSIP | ENSG00000142546 | 0,258804 | 0,032167 |
| PCYOX1L | ENSG00000145882 | -0,2737 | 0,032167 |
| GOLGA8R | ENSG00000186399 | -0,69027 | 0,032167 |
| ZNF444 | ENSG00000167685 | 0,271891 | 0,032168 |
| EFCAB10 | ENSG00000185055 | 0,50222 | 0,032562 |
| ILKAP | ENSG00000132323 | 0,199955 | 0,032677 |
| MRPS18A | ENSG00000096080 | 0,211103 | 0,032792 |
| RNF170 | ENSG00000120925 | -0,34702 | 0,032861 |

| | | | |
|---|---|---|---|
| SDC3 | ENSG00000162512 | -0,20833 | 0,032861 |
| VARS1 | ENSG00000204394 | 0,196714 | 0,032861 |
| RBMS2 | ENSG00000076067 | -0,27463 | 0,032936 |
| ENSG00000273084 | ENSG00000273084 | 0,622851 | 0,032936 |
| IRAK3 | ENSG00000090376 | -0,75995 | 0,033082 |
| ZMIZ1 | ENSG00000108175 | -0,27646 | 0,033082 |
| CDK2AP2 | ENSG00000167797 | 0,298134 | 0,033185 |
| RBM34 | ENSG00000188739 | 0,20858 | 0,033185 |
| SOWAHC | ENSG00000198142 | -0,41945 | 0,033185 |
| EPB41L2 | ENSG00000079819 | -0,18158 | 0,033194 |
| BAIAP2 | ENSG00000175866 | 0,253835 | 0,033294 |
| PTPN13 | ENSG00000163629 | -0,2824 | 0,033307 |
| ERLIN2 | ENSG00000147475 | -0,23561 | 0,033395 |
| NIPAL2 | ENSG00000104361 | -0,43118 | 0,033405 |
| SGPP1 | ENSG00000126821 | -0,32784 | 0,033515 |
| SNHG9 | ENSG00000255198 | 0,393904 | 0,033515 |
| WDR70 | ENSG00000082068 | 0,366005 | 0,033637 |
| C12orf29 | ENSG00000133641 | 0,236367 | 0,033637 |
| ADAMTS1 | ENSG00000154734 | -0,17877 | 0,033637 |
| APOOL | ENSG00000155008 | -0,24828 | 0,033637 |
| SLC35A4 | ENSG00000176087 | 0,178017 | 0,033637 |
| SLC5A3 | ENSG00000198743 | -0,41762 | 0,033637 |
| PCMTD2 | ENSG00000203880 | -0,32197 | 0,033637 |
| RSRC2 | ENSG00000111011 | 0,175241 | 0,033972 |
| DCAF15 | ENSG00000132017 | 0,230412 | 0,033972 |
| EXOSC9 | ENSG00000123737 | 0,212344 | 0,034128 |
| ENSG00000289154 | ENSG00000289154 | 0,441229 | 0,034128 |
| HERC1 | ENSG00000103657 | -0,37428 | 0,034557 |
| EXOSC7 | ENSG00000075914 | 0,220917 | 0,03461 |
| CAMTA2 | ENSG00000108509 | 0,210954 | 0,03461 |
| STAT2 | ENSG00000170581 | -0,25621 | 0,03461 |
| DOHH | ENSG00000129932 | 0,266493 | 0,034711 |
| HSF2BP | ENSG00000160207 | 0,47442 | 0,03477 |
| SNORD3A | ENSG00000263934 | 0,67301 | 0,03477 |
| PER1 | ENSG00000179094 | 0,260269 | 0,03487 |
| IRF2BP1 | ENSG00000170604 | 0,210875 | 0,034906 |
| ASB6 | ENSG00000148331 | 0,194088 | 0,035078 |
| ARMC6 | ENSG00000105676 | 0,180934 | 0,035104 |
| ATP13A1 | ENSG00000105726 | 0,186915 | 0,035183 |
| MRPS14 | ENSG00000120333 | 0,211165 | 0,035228 |
| CAMK4 | ENSG00000152495 | 0,464719 | 0,035256 |
| IMP4 | ENSG00000136718 | 0,179837 | 0,035359 |
| DAAM1 | ENSG00000100592 | -0,35863 | 0,035428 |
| PDE3B | ENSG00000152270 | -0,33873 | 0,035874 |
| SLC30A5 | ENSG00000145740 | -0,24616 | 0,03589 |
| DNAJC30 | ENSG00000176410 | 0,270913 | 0,035965 |
| KAT5 | ENSG00000172977 | 0,232643 | 0,036108 |

| | | | |
|---|---|---|---|
| ASPH | ENSG00000198363 | -0,23979 | 0,036108 |
| DNAJB12 | ENSG00000148719 | 0,183324 | 0,036225 |
| POLE3 | ENSG00000148229 | 0,173735 | 0,036343 |
| NR3C1 | ENSG00000113580 | -0,29333 | 0,036402 |
| IVD | ENSG00000128928 | -0,18949 | 0,036402 |
| EFR3A | ENSG00000132294 | -0,33502 | 0,036402 |
| SF3B4 | ENSG00000143368 | 0,199831 | 0,036402 |
| DNTTIP1 | ENSG00000101457 | 0,213437 | 0,036721 |
| ARHGAP28 | ENSG00000088756 | -0,58236 | 0,036827 |
| DMXL2 | ENSG00000104093 | -0,2687 | 0,036827 |
| CNTNAP3 | ENSG00000106714 | -0,35719 | 0,036827 |
| IMMP1L | ENSG00000148950 | 0,298326 | 0,036827 |
| AP1S2 | ENSG00000182287 | -0,25349 | 0,036827 |
| DUXAP8 | ENSG00000206195 | -0,30597 | 0,036827 |
| ENSG00000280239 | ENSG00000280239 | -0,38416 | 0,036827 |
| SMIM15-AS1 | ENSG00000251279 | 0,656006 | 0,037098 |
| IRF1 | ENSG00000125347 | 0,536292 | 0,037166 |
| NAALAD2 | ENSG00000077616 | -0,62713 | 0,037216 |
| CYTH2 | ENSG00000105443 | 0,202159 | 0,037216 |
| LPGAT1 | ENSG00000123684 | -0,23655 | 0,037216 |
| ST6GALNAC6 | ENSG00000160408 | 0,208256 | 0,037216 |
| H4C15 | ENSG00000270276 | 1,295085 | 0,037216 |
| CBX5 | ENSG00000094916 | -0,21221 | 0,037293 |
| KDM5B | ENSG00000117139 | 0,189232 | 0,037315 |
| COL24A1 | ENSG00000171502 | -0,77047 | 0,037315 |
| ZNF222 | ENSG00000159885 | 0,384969 | 0,037438 |
| C11orf24 | ENSG00000171067 | 0,219628 | 0,037438 |
| ITSN1 | ENSG00000205726 | -0,21919 | 0,037506 |
| PES1 | ENSG00000100029 | 0,213265 | 0,03751 |
| RPA2 | ENSG00000117748 | 0,180532 | 0,037637 |
| FBXL17 | ENSG00000145743 | -0,45055 | 0,037674 |
| GBX2 | ENSG00000168505 | 0,401731 | 0,037674 |
| ENSG00000246465 | ENSG00000246465 | 0,272306 | 0,03775 |
| NBPF15 | ENSG00000266338 | -0,18478 | 0,037827 |
| MECOM | ENSG00000085276 | -0,29087 | 0,03785 |
| RTKN2 | ENSG00000182010 | -0,27442 | 0,03785 |
| ENSG00000279692 | ENSG00000279692 | 0,357644 | 0,03785 |
| PAXBP1-AS1 | ENSG00000238197 | -0,3463 | 0,037872 |
| EGLN1 | ENSG00000135766 | -0,18342 | 0,037979 |
| ENSG00000215014 | ENSG00000215014 | 0,393671 | 0,037979 |
| LONRF2 | ENSG00000170500 | -0,31116 | 0,03801 |
| SPTSSA | ENSG00000165389 | -0,26829 | 0,038083 |
| PAQR8 | ENSG00000170915 | -0,32941 | 0,038083 |
| DDX10 | ENSG00000178105 | 0,204064 | 0,038224 |
| MRPS2 | ENSG00000122140 | 0,201885 | 0,038276 |
| TOX4 | ENSG00000092203 | 0,182268 | 0,038346 |
| TUB | ENSG00000166402 | -0,25031 | 0,038527 |

| | | | |
|---|---|---|---|
| C16orf95-DT | ENSG00000270006 | 0,775983 | 0,038527 |
| PHC3 | ENSG00000173889 | 0,288181 | 0,038606 |
| NUTM2A-AS1 | ENSG00000223482 | 0,216594 | 0,038606 |
| CEBPZ | ENSG00000115816 | 0,208527 | 0,03877 |
| GPD2 | ENSG00000115159 | -0,34056 | 0,039051 |
| RNF32-AS1 | ENSG00000224903 | -0,52486 | 0,039051 |
| ENSG00000262049 | ENSG00000262049 | 0,262507 | 0,039051 |
| ISL2 | ENSG00000159556 | 0,232648 | 0,039423 |
| PARP12 | ENSG00000059378 | -0,72507 | 0,039463 |
| BCS1L | ENSG00000074582 | 0,215256 | 0,039696 |
| ENSG00000275778 | ENSG00000275778 | 0,453317 | 0,039696 |
| TENM1 | ENSG00000009694 | -0,46694 | 0,039704 |
| ENSG00000273010 | ENSG00000273010 | 0,774395 | 0,039974 |
| EIF2B5 | ENSG00000145191 | 0,198205 | 0,040227 |
| NOPCHAP1 | ENSG00000151131 | 0,281521 | 0,040237 |
| CHML | ENSG00000203668 | -0,29156 | 0,040258 |
| KDM7A | ENSG00000006459 | -0,27333 | 0,040435 |
| UTP4 | ENSG00000141076 | 0,166707 | 0,040435 |
| LSM1 | ENSG00000175324 | 0,216168 | 0,040435 |
| FAM215B | ENSG00000232300 | -0,50073 | 0,040435 |
| DXO | ENSG00000204348 | 0,216798 | 0,040544 |
| DPY19L3 | ENSG00000178904 | -0,28674 | 0,040764 |
| PAG1 | ENSG00000076641 | -0,39724 | 0,040845 |
| ENSG00000286786 | ENSG00000286786 | -0,79842 | 0,040902 |
| ZBTB41 | ENSG00000177888 | -0,36403 | 0,040909 |
| CHASERR | ENSG00000272888 | 0,208195 | 0,040978 |
| ADAMTS10 | ENSG00000142303 | -0,40082 | 0,041165 |
| FAM102B | ENSG00000162636 | -0,29377 | 0,041165 |
| RPGRIP1 | ENSG00000092200 | 0,735656 | 0,041214 |
| HM13 | ENSG00000101294 | 0,194722 | 0,04125 |
| SMG9 | ENSG00000105771 | 0,219347 | 0,041602 |
| UBAP1L | ENSG00000246922 | -0,55235 | 0,041643 |
| THUMPD2 | ENSG00000138050 | 0,174561 | 0,041683 |
| COA6-AS1 | ENSG00000231663 | 0,463966 | 0,041683 |
| BCAT1 | ENSG00000060982 | -0,22543 | 0,041688 |
| LMNA | ENSG00000160789 | 0,204091 | 0,041688 |
| NUS1 | ENSG00000153989 | -0,19708 | 0,041841 |
| ZMYND8 | ENSG00000101040 | -0,21108 | 0,042016 |
| USP24 | ENSG00000162402 | -0,22435 | 0,042016 |
| CMTR2 | ENSG00000180917 | -0,34505 | 0,042016 |
| SMN2 | ENSG00000205571 | 0,188293 | 0,042016 |
| ENSG00000277991 | ENSG00000277991 | 0,506068 | 0,042143 |
| SMAD5 | ENSG00000113658 | -0,23357 | 0,042388 |
| COX17 | ENSG00000138495 | 0,238722 | 0,042388 |
| PPAN | ENSG00000130810 | 0,191786 | 0,042481 |
| SURF6 | ENSG00000148296 | 0,201918 | 0,042565 |
| CFAP251 | ENSG00000158023 | 0,66975 | 0,043143 |

| | | | |
|---|---|---|---|
| SOGA1 | ENSG00000149639 | -0,26805 | 0,043482 |
| SPDYE2 | ENSG00000205238 | -0,54768 | 0,043493 |
| ETAA1 | ENSG00000143971 | -0,23925 | 0,044063 |
| KIAA0232 | ENSG00000170871 | -0,23057 | 0,044063 |
| POLR1D | ENSG00000186184 | 0,174663 | 0,044063 |
| KCTD3 | ENSG00000136636 | -0,20015 | 0,044144 |
| POU3F2 | ENSG00000184486 | -0,24125 | 0,044144 |
| SAE1 | ENSG00000142230 | 0,175195 | 0,044201 |
| OTUD6B | ENSG00000155100 | -0,33439 | 0,044388 |
| MLST8 | ENSG00000167965 | 0,188954 | 0,044388 |
| TRMT11 | ENSG00000066651 | 0,203798 | 0,044693 |
| GNB4 | ENSG00000114450 | -0,29268 | 0,044693 |
| SH3BGRL | ENSG00000131171 | -0,22961 | 0,044693 |
| SP2 | ENSG00000167182 | 0,196199 | 0,044693 |
| MECP2 | ENSG00000169057 | 0,179735 | 0,044693 |
| ENSG00000246130 | ENSG00000246130 | 0,31901 | 0,044693 |
| NAB1 | ENSG00000138386 | -0,3463 | 0,044876 |
| ENSG00000269983 | ENSG00000269983 | -0,71483 | 0,044876 |
| MAP1B | ENSG00000131711 | -0,35219 | 0,045085 |
| LANCL1 | ENSG00000115365 | -0,26253 | 0,045089 |
| HSPA1B | ENSG00000204388 | -0,19345 | 0,045089 |
| FAM27E3 | ENSG00000274026 | 0,251182 | 0,045089 |
| PPIE | ENSG00000084072 | 0,205093 | 0,045143 |
| LRFN5 | ENSG00000165379 | -0,54568 | 0,045143 |
| IFRD2 | ENSG00000214706 | 0,186333 | 0,045143 |
| ZSWIM8-AS1 | ENSG00000272589 | 0,477595 | 0,045209 |
| FBXO22 | ENSG00000167196 | 0,249197 | 0,045794 |
| DRG2 | ENSG00000108591 | 0,179901 | 0,045869 |
| ARHGAP29 | ENSG00000137962 | -0,41306 | 0,045869 |
| NCOA2 | ENSG00000140396 | -0,35922 | 0,045869 |
| RNF150 | ENSG00000170153 | -0,32358 | 0,045869 |
| ZNF579 | ENSG00000218891 | 0,244967 | 0,046077 |
| WDR74 | ENSG00000133316 | 0,202163 | 0,046094 |
| ZBTB18 | ENSG00000179456 | -0,2026 | 0,046409 |
| RBM8A | ENSG00000265241 | 0,19163 | 0,046438 |
| DHX30 | ENSG00000132153 | 0,169353 | 0,046622 |
| ZC3H3 | ENSG00000014164 | 0,19849 | 0,046622 |
| MCTS1 | ENSG00000232119 | 0,24627 | 0,046724 |
| GARS1 | ENSG00000106105 | 0,168893 | 0,046889 |
| AKR1A1 | ENSG00000117448 | 0,20534 | 0,046889 |
| SOCS7 | ENSG00000274211 | -0,30021 | 0,046889 |
| FDX2 | ENSG00000267673 | 0,273026 | 0,046918 |
| CCDC90B | ENSG00000137500 | 0,225637 | 0,04709 |
| ZNF516 | ENSG00000101493 | -0,32329 | 0,047171 |
| TRAF3IP1 | ENSG00000204104 | -0,24838 | 0,047184 |
| GAS8 | ENSG00000141013 | 0,187193 | 0,047251 |
| H4C8 | ENSG00000158406 | 0,942485 | 0,047282 |

| | | | |
|---|---|---|---|
| MVP-DT | ENSG00000238045 | 0,211912 | 0,047294 |
| OAZ3 | ENSG00000143450 | 0,486563 | 0,047378 |
| ARC | ENSG00000198576 | 0,533936 | 0,047378 |
| RAB4A-AS1 | ENSG00000177788 | -0,31803 | 0,047475 |
| USP49 | ENSG00000164663 | -0,24973 | 0,047541 |
| SNAPC2 | ENSG00000104976 | 0,264357 | 0,047578 |
| CHAF1A | ENSG00000167670 | 0,206304 | 0,047625 |
| MAD2L1BP | ENSG00000124688 | 0,207886 | 0,047864 |
| ENSG00000263585 | ENSG00000263585 | 0,295836 | 0,047887 |
| PRR3 | ENSG00000204576 | 0,199777 | 0,047941 |
| MANF | ENSG00000145050 | 0,196434 | 0,048085 |
| DSG2 | ENSG00000046604 | -0,26531 | 0,048123 |
| ENSG00000283103 | ENSG00000283103 | 0,295853 | 0,048123 |
| SNRNP40 | ENSG00000060688 | 0,172723 | 0,048173 |
| MAP7D1 | ENSG00000116871 | 0,185746 | 0,048173 |
| GPBP1L1 | ENSG00000159592 | 0,200753 | 0,048173 |
| UBE2J2 | ENSG00000160087 | 0,184856 | 0,048173 |
| GPR153 | ENSG00000158292 | -0,3115 | 0,048208 |
| FOXN3 | ENSG00000053254 | -0,33134 | 0,048377 |
| TRIP6 | ENSG00000087077 | 0,203604 | 0,048377 |
| C1QTNF6 | ENSG00000133466 | -0,31048 | 0,048377 |
| ADAMTS3 | ENSG00000156140 | -0,39498 | 0,048377 |
| TMEM181 | ENSG00000146433 | -0,21239 | 0,04842 |
| MEPCE | ENSG00000146834 | 0,206625 | 0,048482 |
| TRAPPC2B | ENSG00000256060 | 0,378829 | 0,048482 |
| KIAA0586 | ENSG00000100578 | -0,2041 | 0,048507 |
| FBXO30 | ENSG00000118496 | -0,26711 | 0,04877 |
| KRAS | ENSG00000133703 | -0,22047 | 0,048783 |
| RHBDD3 | ENSG00000100263 | 0,212976 | 0,048953 |
| LIPA | ENSG00000107798 | -0,20574 | 0,04905 |
| TSIX | ENSG00000270641 | -0,23546 | 0,049158 |
| PRKCI | ENSG00000163558 | -0,25403 | 0,049162 |
| ZNF766 | ENSG00000196214 | -0,2792 | 0,049162 |
| ASNSP1 | ENSG00000248498 | 0,428686 | 0,049162 |
| MAP3K21 | ENSG00000143674 | -0,22588 | 0,04928 |
| PRKAA1 | ENSG00000132356 | -0,33537 | 0,04938 |
| NECAP2 | ENSG00000157191 | 0,207639 | 0,049506 |
| H2AC6 | ENSG00000180573 | 0,609061 | 0,049506 |
| ENSG00000242588 | ENSG00000242588 | -0,1893 | 0,049506 |
| SLC33A1 | ENSG00000169359 | -0,23535 | 0,049515 |
| ENSG00000288632 | ENSG00000288632 | 0,226051 | 0,04955 |
| RCBTB1 | ENSG00000136144 | -0,26907 | 0,049661 |
| MAPKAPK5-AS1 | ENSG00000234608 | 0,234541 | 0,049725 |
| POU4F1 | ENSG00000152192 | -0,18758 | 0,049809 |
| COPS3 | ENSG00000141030 | 0,168237 | 0,049818 |
| CEP170 | ENSG00000143702 | -0,23801 | 0,049834 |
| ARL4D | ENSG00000175906 | 0,269721 | 0,049834 |

| | | | |
|---|---|---|---|
| KANSL2 | ENSG00000139620 | 0,19399 | 0,04984 |
| LMBRD2 | ENSG00000164187 | -0,31845 | 0,04984 |
