## Supplementary Figure 4 for "Characterization of BoHV-4 ORF45"

**Supplementary Table 4.** Gene Ontology (GO) analysis and Reactome pathway enrichment of annotated genes found to be differentially expressed between treated and Control group (FDR<0.01, LogFC|0.5|).

| Gene Ontology Biological Process | | | |
|---|---|---|---|
| **ID GO:** | **Associated Genes Found** | **Term** | **Pvalue** |
| 51412 | CDKN1A, FOS, FOSL1 | response to corticosterone | 5.7E-04 |
| 35914 | BTG2, FOS, MAFF | skeletal muscle cell differentiation | 2.1E-03 |
| 51385 | CDKN1A, FOS, FOSL1 | response to mineralocorticoid | 2.1E-03 |
| 60389 | GDF15, INHBE, LDLRAD4 | pathway-restricted SMAD protein phosphorylation | 3.4E-03 |
| 44783 | BTG2, CDKN1A, PLK3 | G1 DNA damage checkpoint | 4.8E-03 |
| 31571 | BTG2, CDKN1A, PLK3 | mitotic G1 DNA damage checkpoint | 6.1E-03 |
| 44819 | BTG2, CDKN1A, PLK3 | mitotic G1/S transition checkpoint | 6.1E-03 |
| 42771 | AEN, CDKN1A, PHLDA3 | intrinsic apoptotic signaling pathway in response to DNA damage by p53 class mediator | 6.4E-03 |
| 72413 | BTG2, CDKN1A, PLK3 | signal transduction involved in mitotic cell cycle checkpoint | 7.0E-03 |
| 1902402 | BTG2, CDKN1A, PLK3 | signal transduction involved in mitotic DNA damage checkpoint | 7.0E-03 |
| 1902403 | BTG2, CDKN1A, PLK3 | signal transduction involved in mitotic DNA integrity checkpoint | 7.0E-03 |
| 60393 | GDF15, INHBE, LDLRAD4 | regulation of pathway-restricted SMAD protein phosphorylation | 7.3E-03 |
| 72431 | BTG2, CDKN1A, PLK3 | signal transduction involved in mitotic G1 DNA damage checkpoint | 7.7E-03 |
| 1902400 | BTG2, CDKN1A, PLK3 | intracellular signal transduction involved in G1 DNA damage checkpoint | 7.7E-03 |
| 6977 | BTG2, CDKN1A, PLK3 | DNA damage response, signal transduction by p53 class mediator resulting in cell cycle arrest | 8.4E-03 |
| **Reactome Pathway** | | | |
| **ID R-HSA:** | **Associated Genes Found** | **Term** | **Pvalue** |
| 2559586 | CDKN1A, H1-2, H2AC19, H2BC11, H2BU1, H4C11, H4C14 | DNA Damage/Telomere Stress Induced Senescence | 4.7E-07 |
| 2559582 | CDKN1A, FOS, H2AC19, H2BC11, H2BU1, H4C11, H4C14 | Senescence-Associated Secretory Phenotype (SASP) | 4.6E-06 |
| 2559583 | CDKN1A, FOS, H1-2, H2AC19, H2BC11, H2BU1, H4C11, H4C14 | Cellular Senescence | 1.6E-05 |
| 9616222 | CDKN1A, H2AC19, H2BC11, H2BU1, H4C11, H4C14 | Transcriptional regulation of granulopoiesis | 2.8E-05 |
| 3214815 | H2AC19, H2AW, H2BC11, H2BU1, H4C11, H4C14 | HDACs deacetylate histones | 3.5E-05 |
| 171306 | H2AC19, H2BC11, H2BU1, H4C11, H4C14 | Packaging Of Telomere Ends | 3.9E-05 |
| 110330 | H2AC19, H2BC11, H2BU1, H4C11, H4C14 | Recognition and association of DNA glycosylase with site containing an affected purine | 5.5E-05 |
| 110331 | H2AC19, H2BC11, H2BU1, H4C11, H4C14 | Cleavage of the damaged purine | 5.5E-05 |
| 73927 | H2AC19, H2BC11, H2BU1, H4C11, H4C14 | Depurination | 5.5E-05 |
| 110328 | H2AC19, H2BC11, H2BU1, H4C11, H4C14 | Recognition and association of DNA glycosylase with site containing an affected pyrimidine | 8.2E-05 |
| 110329 | H2AC19, H2BC11, H2BU1, H4C11, H4C14 | Cleavage of the damaged pyrimidine | 8.2E-05 |

| | | | |
|---|---|---|---|
| 73928 | H2AC19, H2BC11, H2BU1, H4C11, H4C14 | Depyrimidination | 8.2E-05 |
| 73728 | H2AC19, H2BC11, H2BU1, H4C11, H4C14 | RNA Polymerase I Promoter Opening | 9.4E-05 |
| 73929 | H2AC19, H2BC11, H2BU1, H4C11, H4C14 | Base-Excision Repair. AP Site Formation | 9.4E-05 |
| 5334118 | H2AC19, H2BC11, H2BU1, H4C11, H4C14 | DNA methylation | 1.1E-04 |
| 5625886 | H2AC19, H2BC11, H2BU1, H4C11, H4C14 | Activated PKN1 stimulates transcription of AR (androgen receptor) regulated genes KLK2 and KLK3 | 1.2E-04 |
| 427359 | H2AC19, H2BC11, H2BU1, H4C11, H4C14 | SIRT1 negatively regulates rRNA expression | 1.3E-04 |
| 2559580 | FOS, H2AC19, H2BC11, H2BU1, H4C11, H4C14 | Oxidative Stress Induced Senescence | 1.6E-04 |
| 212300 | H2AC19, H2BC11, H2BU1, H4C11, H4C14 | PRC2 methylates histones and DNA | 1.7E-04 |
| 2299718 | H2AC19, H2BC11, H2BU1, H4C11, H4C14 | Condensation of Prophase Chromosomes | 1.7E-04 |
| 606279 | H2AC19, H2BC11, H2BU1, H4C11, H4C14 | Deposition of new CENPA-containing nucleosomes at the centromere | 1.7E-04 |
| 774815 | H2AC19, H2BC11, H2BU1, H4C11, H4C14 | Nucleosome assembly | 1.7E-04 |
| 427389 | H2AC19, H2BC11, H2BU1, H4C11, H4C14 | ERCC6 (CSB) and EHMT2 (G9a) positively regulate rRNA expression | 2.0E-04 |
| 1221632 | H2AC19, H2BC11, H2BU1, H4C11, H4C14 | Meiotic synapsis | 2.2E-04 |
| 3214847 | H2AC19, H2AW, H2BC11, H2BU1, H4C11, H4C14 | HATs acetylate histones | 2.6E-04 |
| 912446 | H2AC19, H2BC11, H2BU1, H4C11, H4C14 | Meiotic recombination | 3.1E-04 |
| 9018519 | FOS, H2AC19, H2BC11, H2BU1, H4C11, H4C14 | Estrogen-dependent gene expression | 3.3E-04 |
| 157579 | H2AC19, H2BC11, H2BU1, H4C11, H4C14 | Telomere Maintenance | 3.6E-04 |
| 201722 | H2AC19, H2BC11, H2BU1, H4C11, H4C14 | Formation of the beta-catenin:TCF transactivating complex | 3.6E-04 |
| 5250924 | H2AC19, H2BC11, H2BU1, H4C11, H4C14 | B-WICH complex positively regulates rRNA expression | 3.6E-04 |
| 73772 | H2AC19, H2BC11, H2BU1, H4C11, H4C14 | RNA Polymerase I Promoter Escape | 3.6E-04 |
| 73884 | H2AC19, H2BC11, H2BU1, H4C11, H4C14 | Base Excision Repair | 3.7E-04 |
| 1912408 | H2AC19, H2BC11, H2BU1, H4C11, H4C14 | Pre-NOTCH Transcription and Translation | 3.7E-04 |
| 5625740 | H2AC19, H2BC11, H2BU1, H4C11, H4C14 | RHO GTPases activate PKNs | 3.9E-04 |
| 8936459 | H2AC19, H2BC11, H2BU1, H4C11, H4C14 | RUNX1 regulates genes involved in megakaryocyte differentiation and platelet function | 4.1E-04 |
| 5250913 | H2AC19, H2BC11, H2BU1, H4C11, H4C14 | Positive epigenetic regulation of rRNA expression | 5.8E-04 |
| 5250941 | H2AC19, H2BC11, H2BU1, H4C11, H4C14 | Negative epigenetic regulation of rRNA expression | 5.9E-04 |
| 5578749 | H2AC19, H2BC11, H2BU1, H4C11, H4C14 | Transcriptional regulation by small RNAs | 5.9E-04 |
| 73864 | H2AC19, H2BC11, H2BU1, H4C11, H4C14 | RNA Polymerase I Transcription | 5.9E-04 |
| 1912422 | H2AC19, H2BC11, H2BU1, H4C11, H4C14 | Pre-NOTCH Expression and Processing | 5.9E-04 |
| 427413 | H2AC19, H2BC11, H2BU1, H4C11, H4C14 | NoRC negatively regulates rRNA expression | 5.9E-04 |
| 977225 | H2AC19, H2BC11, H2BU1, H4C11, H4C14 | Amyloid fiber formation | 5.9E-04 |
| 73854 | H2AC19, H2BC11, H2BU1, H4C11, H4C14 | RNA Polymerase I Promoter Clearance | 6.1E-04 |
| 1500620 | H2AC19, H2BC11, H2BU1, H4C11, H4C14 | Meiosis | 6.4E-04 |
| 73886 | H2AC19, H2BC11, H2BU1, H4C11, H4C14 | Chromosome Maintenance | 6.4E-04 |
| 5617472 | H2AC19, H2BC11, H2BU1, H4C11, H4C14 | Activation of anterior HOX genes in hindbrain development during early embryogenesis | 6.5E-04 |
| 5619507 | H2AC19, H2BC11, H2BU1, H4C11, H4C14 | Activation of HOX genes during differentiation | 6.5E-04 |
| 5693571 | H2BC11, H2BU1, H4C11, H4C14 | Nonhomologous End-Joining (NHEJ) | 9.1E-04 |
| 3214858 | H2AC19, H2AW, H4C11, H4C14 | RMTs methylate histone arginines | 1.0E-03 |

| | | | |
|---|---|---|---|
| 5693606 | H2BC11, H2BU1, H4C11, H4C14 | DNA Double Strand Break Response | 1.1E-03 |
| 5693607 | H2BC11, H2BU1, H4C11, H4C14 | Processing of DNA double-strand break ends | 1.2E-03 |
| 5693565 | H2BC11, H2BU1, H4C11, H4C14 | Recruitment and ATM-mediated phosphorylation of repair and signaling proteins at DNA double strand breaks | 1.2E-03 |
| 6791312 | BTG2, CDKN1A, PLK3 | TP53 Regulates Transcription of Cell Cycle Genes | 1.3E-03 |
| 198725 | FOS, FOSL1, TPH1 | Nuclear Events (kinase and transcription factor activation) | 1.3E-03 |
| 9031628 | FOS, FOSL1, TPH1 | NGF-stimulated transcription | 1.4E-03 |
| 69473 | H2BC11, H2BU1, H4C11, H4C14 | G2/M DNA damage checkpoint | 1.7E-03 |

PValue was corrected with Bonferroni step down
